# Quantitative and population genomics suggest a broad role of staygreen loci in the drought adaptation of sorghum

**DOI:** 10.1101/2021.06.09.447769

**Authors:** Jacques M. Faye, Eyanawa A. Akata, Bassirou Sine, Cyril Diatta, Ndiaga Cisse, Daniel Fonceka, Geoffrey P. Morris

**Author notes:** Author for correspondence: Geoffrey P. Morris.

## Abstract

- Drought is a major constraint on plant productivity globally. Sorghum (*Sorghum bicolor*) landraces have evolved in drought-prone regions, but the genetics of their adaptation is not yet understood. Loci underlying stay-green post-flowering drought tolerance (*Stg*), have been identified in a temperate breeding line, but their role in drought adaptation of tropical sorghum is to be elucidated.
- We phenotyped 590 diverse sorghum accessions from West Africa under field-based managed drought stress, pre-flowering (WS1) and post-flowering (WS2) over several years and conducted genome-wide association studies (GWAS).
- Broad-sense heritability for grain and biomass yield components was high (33-92%) across environments. There was a significant correlation between stress tolerance index (STI) for grain weight across WS1 and WS2. GWAS revealed that *SbZfl1* and *SbCN12*, orthologs of maize flowering genes, likely underlie flowering time variation under these conditions. GWAS further identified associations (n = 134) for STI and drought effects on yield components, including 16 putative pleiotropic associations. Thirty of the associations colocalized with *Stg1–4* loci and had large effects. Seven lead associations, including some within *Stg1*, overlapped with positive selection outliers.
- Our findings reveal natural genetic variation for drought tolerance-related traits, and suggest a broad role of *Stg* loci in drought adaptation of sorghum.

## INTRODUCTION

Unpredictable rainfall and drought are major limitations to plant productivity worldwide. Improving crop adaptation to water limitation is critical for establishing food security in developing countries where smallholder farmers are vulnerable to climate change (Mundia *et al*., 2019). From an agronomic perspective, drought adaptation is the ability to maintain yield under agronomic water limitation (Blum, 2010). An understanding of the genetic architecture of grain yield and its components across various drought scenarios can facilitate crop breeding to increase production. However, collecting good phenotypic data under well-managed water stress environments and integrating phenotypes with genotypes remain major constraints. The genetic dissection of yield components under various drought scenarios would provide favorable natural variants for drought tolerance.

Sorghum (*Sorghum bicolor*) is a staple cereal crop in drought-prone regions worldwide, including many developing countries of the semi-arid tropics as well as industrialized countries in the temperate latitudes. Sorghum is among the most drought-resilient crops, but the physiological and genetic basis of its drought tolerance is not yet understood (Mullet *et al*., 2014). Several quantitative trait loci (QTL) associated with drought tolerance variation in sorghum have been identified, but no genes have been cloned. The best studied of these QTL are stay-green loci (*Stg1–Stg4)* linked to post-flowering drought tolerance in biparental families and near-isogenic lines (Tuinstra *et al*., 1997; Xu *et al*., 2000; Harris *et al*., 2007; Borrell *et al*., 2014b; Hayes *et al*., 2016). The *Stg* loci influence several aspects of sorghum development, including canopy architecture, water use, and grain yield (Borrell *et al*., 2014b). *Stg* alleles were identified in a temperate-adapted breeding line BTx642 (formerly B35) that is derived from a tropically-adapted Ethiopian durra landrace (IS12555). However, the prevalence of the *Stg* alleles in sub-Saharan Africa or their role in drought adaptation (if any) is not known. Understanding the genetic basis of drought adaptation in sorghum could elucidate the process of environmental adaptation and facilitate breeding of drought-tolerant varieties.

Local varieties have been under natural and farmers selection for adaptation to environmental conditions and farming systems. Local varieties of sorghum have adapted to various environmental conditions since their domestication (Harlan & De Wet, 1972; Wendorf *et al*., 1992). Consequently, positive pleiotropic loci for combined pre- and post-flowering drought tolerance might exist in locally-adapted varieties. West African sorghum is extremely diverse and there have been few cycles of selection in breeding programs (Mauboussin *et al*., 1977; Leiser *et al*., 2014). The West African sorghum association panel (WASAP), including landraces and breeding lines that consist of working collections of breeding programs, was assembled and genotyped using genotyping-by-sequencing technology. However, the genetic architecture underlying grain yield and its components under various drought scenarios remains largely unknown in the germplasm. We hypothesized that positively pleiotropic QTLs confer combined pre- and post-flowering drought tolerance in the West African sorghum.

Genome-wide association studies (GWAS) contribute to the identification of natural variants, taking advantage of historical recombinations within diversity panels (Yu & Buckler, 2006; McCouch *et al*., 2016; Yano *et al*., 2016; Zhao *et al*., 2019). A grass species such as sorghum is suitable to identify natural variants underlying complex agronomic traits partly due to its small genome size and moderate LD (Paterson *et al*., 2009; Mace *et al*., 2013; McCormick *et al*., 2018). Disentangling positive pleiotropic effects of drought-yield QTLs through GWAS can contribute to detect and characterize the natural allelic variation existing within locally-adapted populations. In this study, we performed GWAS on 756 sorghum accessions of the WASAP under ten different environments using the previous GBS SNP dataset. We (*i*) characterize the genetic variation of yield components under various water stress environments; (*ii*) identify genetic variants at known and novel drought tolerance loci with high productivity under pre- and post-flowering water stress; (*iii*) investigate the pleiotropic effect of drought tolerance QTLs associated with STI and reduction of yield components under various drought scenarios; and (*iv*) determine signatures of selection overlapping with identified drought tolerance QTLs. The present study provides knowledge of the genetic architecture of yield components under various drought scenarios.

## MATERIALS AND METHODS

### Plant materials

The West African Sorghum Association Panel (WASAP) consists of *N* = 756 genotyped accessions from the four West African countries of Senegal (118 accessions), Mali (123), Togo (156), and Niger (359) (Faye *et al*., 2021) (Fig. 1a). The panel includes predominantly landraces along with some local breeding lines and local improved varieties. Five local breeding lines were used as checks for use in augmented design: T1 (IRAT 204/CE151-262), T2 (CE145-266), T3 (ISRA-621B/Faourou), T4 (CE180-33), and T5 (53-49). Two international drought-response reference lines, Tx7000 (pre-flowering drought tolerant, post-flowering drought susceptible) and BTx642 (pre-flowering drought susceptible, post-flowering drought tolerant), were used as controls (Burke *et al*., 2013; Borrell *et al*., 2014a).

**Fig. 1.**
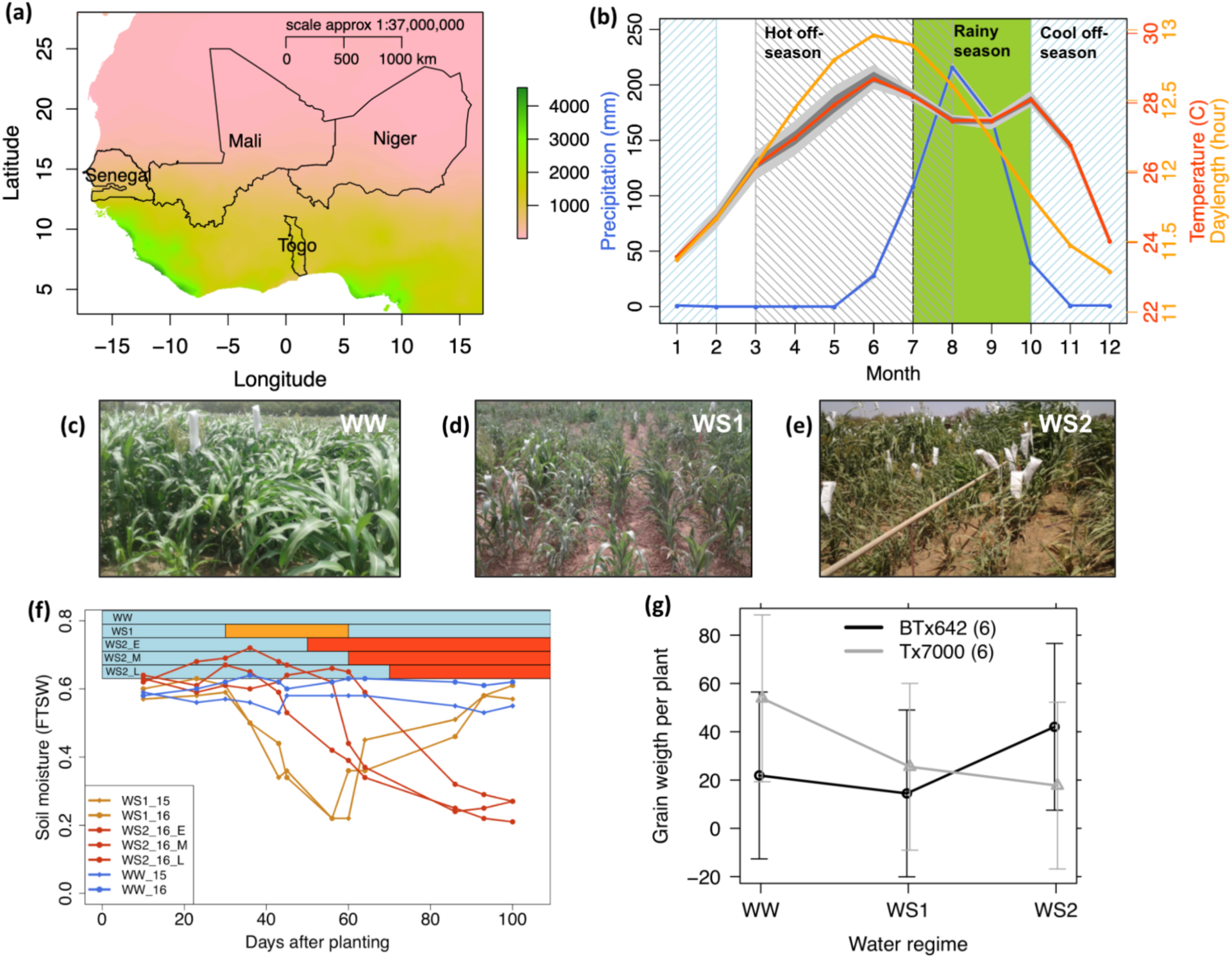
Experimental system to study drought stress response of diverse sorghum germplasm. (a) The four countries of origin for sorghum accessions in the West Africa Sorghum Association Panel (WASAP) with the West African precipitation gradient noted by the color scale. (b) Average monthly precipitation, temperature, and daylength at the experimental station in Bambey, Senegal. The green block represents the rainy season (“hivernage”) when farmers grow crops and when we conducted rainfed experiments. The gray-striped block indicates the hot off-season and the blue-striped block indicates the cool off-season when we conducted managed drought stress. (c-e) Photos of plants under (c) well-watered (WW), (d) pre-flowering water stress (WS1), and (e) post-flowering water stress (WS2) environments. (f) Fraction of transpirable soil water in WW (blue lines), WS1 (orange lines), and WS2 (red lines) during 2015 (line with diamond shape dots) and 2016 (line with close circle dots) off-seasons. Horizontal bars indicate the water stress application periods for WS1 (orange) and WS2 (red) relative to WW (light blue). The three red bars/lines for WS2 represent three maturity groups (E: early maturity, M: medium maturity, L: late maturity) that the panel was divided into so that post-flowering water stress could be applied consistently relative to flowering. (g) Cross genotype x environment interaction of the pre-flowering (Tx7000) and post-flowering (BTx642) drought tolerance checks across WW, WS1, and WS2 environments. The error bars represent 95% confidence intervals.

### Field trials

Field experiments were performed over four years (2014–2017) in Senegal at the Bambey Research Station, CNRA–Centre National de Recherche Agronomique (14.42°N, 16.28°W) in the Soudano-Sahelian zone (Fig. 1a). The average annual precipitation is ∼600 m, which occurs strictly in the rainy season (“hivernage”) of July to October, with maximum monthly precipitation typically occurring in August (Fig. 1b). In total, ten experiments were performed in an incomplete randomized block design (augmented block design) across the four years (Table 1; Fig. S1a-f). The experimental set-up followed a column–row field layout with 30 blocks for 2014 experiments or 25 blocks for 2015-2017 experiments, with 19 genotypes and the 5 local check varieties (present in each block for spatial variation analysis) within each block. Each entry was sown in a 3 m row with 0.6 m space between rows and 0.2 m space between plants (or hills) within a row. Each entry was surrounded by one row of fill material (IRAT 2014). Ten days after planting, plants were thinned to keep only one plant per hill, for a density of about 84,000 plants ha^-1^. Two experiments were carried out under rainfed conditions (RF) during the rainy season in 2014 with one-month planting date interval: RF1 (planted in August) and RF2 (planted in September). Managed drought stress experiments were conducted in the off-season to take advantage of the complete lack of precipitation during the Sahelian dry season (Fig. 1b).

**Table 1.** Details of field experiments.

| Treatment <sup>a</sup> | Season | Year <sup>b</sup> | Period | Trial code |
| --- | --- | --- | --- | --- |
| Rainfed | Rainy season | 2014 | July–October | RF1 |
| Rainfed | Rainy season | 2014 | July–October | RF2 |
| Well-watered | Hot off-season | 2015 | March–August | WW_15 |
| Pre-flowering | Hot off-season | 2015 | March–August | WS1_15 |
| Well-watered | Cool off-season | 2016 | October–February | WW_16 |
| Pre-flowering | Cool off-season | 2016 | October–February | WS1_16 |
| Post-flowering | Cool off-season | 2016 | October–February | WS2_16 |
| Well-watered | Cool off-season | 2017 | October–February | WW_17 |
| Pre-flowering | Cool off-season | 2017 | October–February | WS1_17 |
| Post-flowering | Cool off-season | 2017 | October–February | WS2_17 |

<sup>a</sup> Pre-flowering and post-flowering denote the pre- or post-flowering water stress environments;
<sup>b</sup> The year where the majority of the experiment took place; RF, rainfed condition; WW, well water; WS1, pre-flowering water stress; WS2, post-flowering water stress.

### Managed drought stress

Well-watered (WW) and pre-flowering water stress (WS1) experiments were planted during the hot off-season in 2015 (March to August). Three experiments, under WW, WS1, and post-flowering water stress (WS2), were planted during the cool off-season in 2015–2016 (October 2015 to March 2016; note as “2016” experiments) and 2016-2017 (October 2016 to March 2017; noted as “2017” experiments). During the rainy season of 2014, the cumulative rainfall recorded was 395 mm. The average daily temperature ranges between 22.4 and 35 °C and average relative humidity between 42 and 89%. For WW, irrigation was applied twice a week (30 mm each time) until physiological maturity. For WS1, water limitation was applied 30 DAP, to mimic a one-month pre-flowering drought, and irrigation was restarted 60 DAP until physiological maturity. For the WS2, water limitation was applied when 75% of plants in a maturity group flowered and maintained until physiological maturity. Three maturity groups were defined based on accession phenology characterized during 2014 experiments for water deficit application in WS2. The fraction of transpirable soil water (FTSW) in different managed drought stress experiments was determined using a DIVINER 2000 (Sentek Pty Ltd, Adelaide, SA).

### Phenotypic measurements

In each environment, phenological, physiological, and yield component traits were measured. Days to 50% flowering (DFLo) of plants in a plot (one row), above-ground dry biomass (DBM), plant height (PH), and yield components including grain weight per panicle (GrW), panicle weight (PW), grain number per plant (GrN), and thousand-grain weight (TGrW) were measured and used for association mapping studies. For each trait except for DFLo and TGrW, three plants from the middle row of each plot were used for measurements. The drought stress tolerance index (STI) (Li *et al*., 2018a; Yuan *et al*., 2019) for grain weight was calculated from the GrW under WW and WS1 or WS2 as follows:

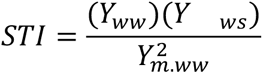

Where *Y_ww_* and *Y_ws_* is the grain weight of a given genotype in WW and WS environments, respectively, and *Y_m.ww_* is the mean value of GrW in the WW environment. For the STI, the higher the value, the more tolerant the genotype to the stress. The drought reduction of each yield component relative to the control environment was calculated as follow:

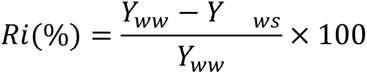

Where *Ri* is the drought response of a genotype for trait *i*, *Y_ww_* and *Y_ws_* are the performance of the genotype in the control environment and water-stressed environment, respectively.

### Statistical analysis of phenotypes

Each year-treatment combination is considered an environment. Statistical analysis was performed using the R program (R Core Team, 2016). Spatial variation within each environment was analyzed based on the check varieties in each block using the *SpATS* package (Rodríguez-Álvarez *et al*., 2018) to obtain genotype-adjusted means. The variance components were estimated by fitting the mixed linear model with random effects for all genotypes (G), water regimes (WR), years (Y), and GxY interaction effects using the *lme4* package (Bates et 2010). Broad sense heritability (*H^2^*) was calculated based on variance components derived from the mixed effect model. *H^2^* was estimated for each trait across environments based on the genotypic variance and the total phenotypic variance. Phenotypic correlations among traits were calculated using the Pearson correlation coefficient of the PerformanceAnalytics package (Peterson *et al*., 2014). Tukey’s Honestly Significant Difference (TukeyHSD) test in the Agricolae package (Mendiburu, 2009) was used to test the difference of genotype performance between environments or botanical types. The BLUP values of the phenotypes were calculated by combining data for a given water regime across years or across all environments. The phenotypic BLUPs and genotype-adjusted means were used for the genome-wide association analysis across environments.

### Genome-wide association studies

To identify drought-yield QTLs, GWAS was performed using the general linear model (GLM) with principal component (PC) eigenvalues and mixed linear model (MLM) in the GAPIT package (Lipka *et al*., 2012). These two GWAS models were used as complementary because the GLM may identify false-positive associations while MLM may lead to false-negative associations when controlling for false-positive significant associations. The SNP dataset was filtered for MAF > 0.02, which corresponds to >15 observations of the minor allele within the panel of *N* = 756 genotyped accessions. The first five PCs and the kinship matrix were used to account for population structure and genetic relatedness effects, respectively for the MLM. The significance level of GWAS associations were defined based on Bonferroni-corrected *p-*value 0.05 for the GLM with PC (referred to as GLM+Q hereafter) or at least top five SNPs above *p* < 10^-5^ cutoff for the MLM. The most highly-associated SNP (“lead SNP”) within a 150 kb genomic region defined based on average linkage disequilibrium (LD) decay in global sorghum germplasm (Morris *et al*., 2013) was chosen to represent the association. A list of a priori candidate genes of cloned cereal flowering times from a previous study (Faye *et al*., 2019) was used for colocalization analysis between lead SNP and candidate genes.

### Locus-specific analyses

LD heatmaps were constructed using the R package *LD heatmap 0.99-4* (Shin *et al*., 2006). BLUP values of phenotypes across water stress environments were used for the estimation of the proportion of phenotypic variance explained (PVE) by lead SNPs from the GWAS. The PVE was estimated using linear models with fractions of ancestry inferred by ADMIXTURE (Alexander *et al*., 2009) used as fixed covariates. Statistical enrichment analysis for colocalization between GWAS lead SNPs and all *Stg* QTLs from the sorghum QTL Atlas (Mace *et al*., 2019) was performed based on 1000 permutation tests. Statistical significance was assessed with a two-sample *t*-test with α = 0.05. Geographic distribution of the associated lead SNP alleles with DFLo or putative drought tolerance was determined using an existing set of georeferenced global sorghum landraces (Lasky *et al*., 2015). Lead associations within *Stg1-3* QTLs were selected based on their association with drought tolerance variables, LD with other lead associations within a locus, contribution to the phenotypic variation, and availability in the GBS data for global sorghum landraces.

### Genome-wide selection scans

For selection scans, we included 550 worldwide sorghum accessions including wild relative sorghum accessions with available sequencing data (Morris *et al*., 2013). Genome-wide selection scans were performed based on 100 kb sliding windows using the vcftools program (Danecek *et al*., 2011). Decreased genome-wide nucleotide diversity (π) in durra-caudatum, durra, and guinea cultivars relative to wild relatives was performed to assess domestication and diversification selections for drought responses to dry (in durra-caudatum and durra genome) versus humid (in guinea genome) regions. Statistical enrichment analysis for colocalization between *π* outlier regions and *Stg1–4* loci was performed based on 1000 permutation tests. Statistical significance of mean differences were based on two-sided two-sample *t*-tests with α = 0.05.

## RESULTS

### Phenotypic variation for drought tolerance related traits

A total of 590 WASAP accessions were evaluated for phenological, physiological, and yield component traits under ten environments across four years in Senegal (Fig. 1a,b; Fig. S1). To assess the level of drought stress applied, we estimated the fraction of transpirable soil water (FTSW) in the WW, WS1, and WS2 (Fig. 1c-f). FTSW was estimated to be 0.6 in both WW and stressed treatment before water deficit treatment, then dropped to ∼0.2 and 0.3 in WS1 and WS2 environments, respectively. To assess the effect of each water condition, we characterized the grain yield components and days to flowering of genotypes. A non-significant cross-over genotype-environment interaction (*p* < 0.08) was observed between the two drought tolerance reference lines, BTx642 and Tx7000 in WS1 and WS2 (Fig. 1g). As expected, the average grain weight and number of genotypes was significantly reduced in WS1 and WS2 relative to WW treatment (Fig. S1a,b). Overall, DFLo was significantly delayed in 2015 hot off-season environments, whereas it was reduced in cool off-seasons of 2016 and 2017 relative to rainfed conditions (Fig. S1c). DFLo was delayed in WS1, whereas it was not different in WS2 relative to the WW controls. DBM was significantly reduced in all stressed environments, except in WS1 of 2015 relative to RF (Fig. S1d). Average grain weight was not significantly different between RF and WS2 (Fig. S1e). The average GrN was significantly lower in WS1 than in WS2 (Fig. S1f).

### Genetic variation in drought response

Broad-sense heritability (*H*^2^) estimates varied from moderate to high with values ranging from 33% for GrN to 92% for PH in the whole WASAP (Table S1). The average grain weight was not significantly different between caudatum accessions and durra and guinea accessions within each water regime in terms of production under drought stress (Fig. 2a). The durra-caudatum intermediates had significantly higher average grain weight than caudatum (13%, *p* < 0.05) and guinea (16%, *p* < 0.05) accessions, but not with durra (7%, *p* < 0.1) accessions. The average GrN was not significantly different between botanical types (Fig. 2b). Significant correlations were observed among yield components, including GrW, DBM, and STI for grain weight, across WS1 and WS2 regimes (Fig. S2a). High positive correlation was observed between BLUP of GrW, PW, DBM, and GrN, while TGrW was negatively correlated with grain number (Fig. S2b). Significant correlations were observed between DBM in WS1 and WS2 and other yield components, GrW, GrN, DFLo, and PH across RF conditions (Fig. S2c,d). Overall, genetic differences contributed to the phenotypic variation in managed water stress conditions.

**Fig. 2.**
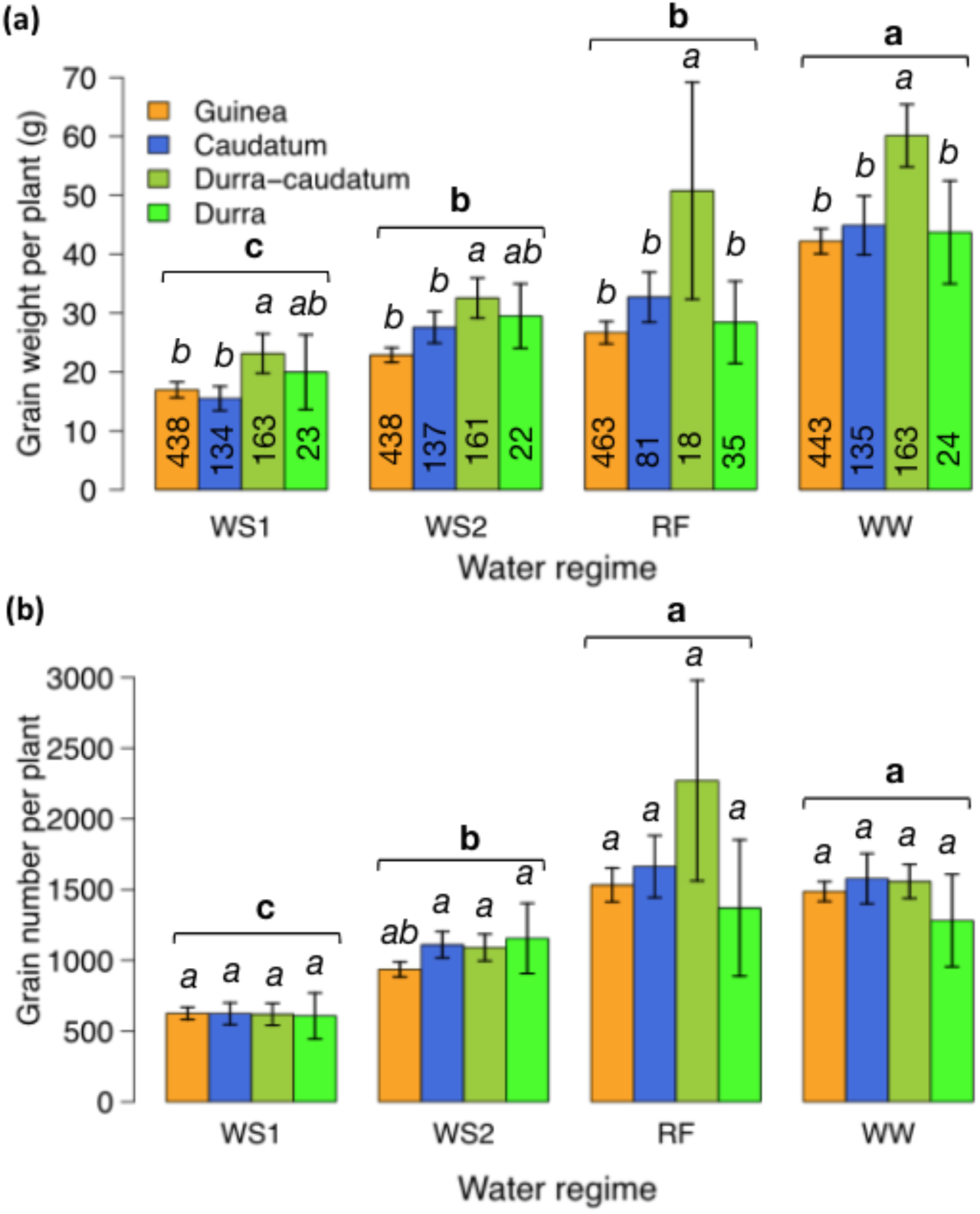
Effects of managed drought stress on grain yield components. (a) Differences in grain weight among botanical types within each water regime, rainfed condition (RF), well-watered (WW), pre-flowering water stress (WS1), post-flowering water stress (WS2). Numbers within bar plots indicate the number of genotypes per botanical type in each water regime (two environments in each). (b) Differences in grain number among botanical types within each water regime. The letters indicate Tukey honestly statistical difference at α = 0.05, with bold letters indicating the across water regime comparison and italics letters representing the across botanical type comparison within the water regime.

### Genome-wide association studies of flowering time

To identify loci potentially underlying quantitative trait variation in West African sorghum, we carried out GWAS using 130,709 SNP markers. First, we considered DFLo under WW off-season environments of 2015, 2016, and 2017 and BLUPs across all off-season environments to map known flowering time candidate genes using GLM+Q. No significant peak above the Bonferroni-corrected *p*-value of 0.05 was identified for DFLo of the 2015 data, but significant associations were identified for DFLo of the 2016 and 2017 data (Fig. S3). Two SNPs, S6_55280640 and S3_62811196, were significantly associated with DFLo in both years, and co-localized with *a priori* candidate flowering time genes *SbZfl1* (Sobic.006G201600; 9 kb away) and *SbCN12* (Sobic.003G295300; 61 kb away), respectively. In both 2016 and 2017, S6_55280640 was the lead SNP (*p* < 10^-10^ in 2016; *p* < 10^-10^ in 2017) of the associated region on chromosome 6. A third SNP, S2_67812515, was significantly associated with DFLo in 2017 data and colocalized with the *a priori* candidate gene *Maturity2* (Sobic.002G302700; 70 kb away). Significant associations were not identified above the Bonferroni threshold (*p* > 10^-5^) when the MLM with PCA and kinship matrix were used to account for both population structure and genetic relatedness effects (Fig. S3).

The same associated SNPs near *SbZfl1* and *SbCN12*, noted above, were observed for flowering time BLUPs across all off-season environments (Fig. 3a; File S1). Lead SNP S6_55280640 was located one gene away from *SbZfl1* (Fig. 3c). The T allele of S6_55280640, associated with shorter flowering times under RF conditions (Fig. 3d), had a wide geographic distribution and was found at high frequency in accessions of the Sahel, Ethiopia, and west India (Fig. 3e). Lead SNP S3_62811196 was the top association near *SbCN12* (Fig. 3f). The T allele of S3_62811196, associated with short flowering times under RF conditions (Fig. 3g), is globally-rare, found mostly in accessions from Niger and northern Nigeria (Fig. 3h).

**Fig. 3.**
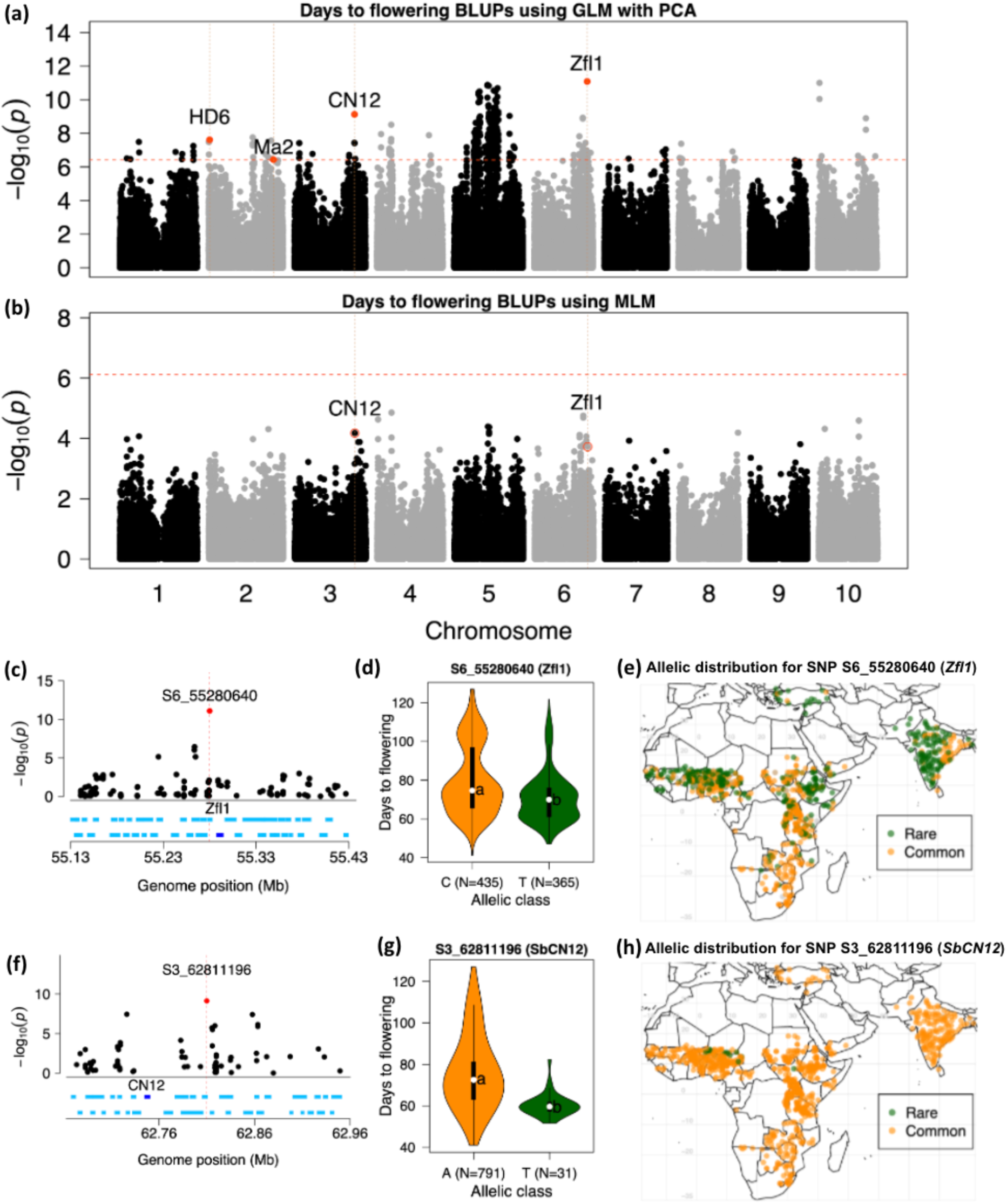
Genome-wide associations for flowering time. (a, b) Manhattan plots for days to flowering (DFLo) for best linear unbiased predictor (BLUPs) across all off-season environments over three years using (a) general linear model with principal components (GLM+Q) and (b) mixed-linear model (MLM). Horizontal dashed line indicates the Bonferroni correction at 0.05. Red dots indicate peak SNPs colocalizing (based on 150 kb cutoff) with *a priori* candidate genes for flowering time. (c) Zoomed-in Manhattan plot for the GLM+Q of a 150 kb region on chromosome 6 around the lead associated SNP, S6_55280640 that colocalizes with *a priori* candidate gene *Zfl1* (dark blue segment). (d) Days to flowering across rainfed environments by allelic classes of S6_55280640 in the WASAP. Letters within violin pots indicate Tukey’s honestly significant difference at α = 0.05. (e) Geographic distribution of early (T) and late (C) flowering-associated alleles of S6_55280640 in global sorghum landraces. (f) Zoomed-in Manhattan plot for the GLM+Q of a 150 kb region on chromosome 3 around the lead associated SNP, S3_62811196 that colocalizes with *a priori* candidate gene *SbCN12* (dark blue segment). (g) Days to flowering across rainfed environments by allelic classes of S3_62811196 in the WASAP. (h) Geographic distribution of the early (T) and late (A) flowering-associated alleles of S3_62811196 in global sorghum landraces.

### Genome-wide association studies for drought tolerance

To generate hypotheses on the loci that underlie drought tolerance variation in sorghum, we performed GWAS for grain weight STI and the reduction of PW (RPW), DBM (RDBM), GrN (RGrN), PH (RPH), TGrW (RTGrW) in water-stressed environments. We considered water-stress scenarios separately (WW vs. WS1, WW vs. WS2) and together (WW vs. WS1 and WS2). In total, 222 and 214 associations were identified by the GLM+Q and MLM, respectively for drought response variables and STI in all drought stress environments (File S2; Fig. S4). Among the associations, 134 were commonly identified by both GWAS methods.

To determine QTLs that have positive pleiotropic effect on pre- and post-flowering drought tolerance among the associations above, we looked for common associations across different water-stressed environments. We defined a pleiotropic QTL as one lead SNP or locus being mapped in both pre- and post-flowering drought scenarios, or associated with several drought response variables. Among the associations, 16 putative pleiotropic associations for drought responses were observed across water stress environments (Table S2). For example, the SNP S4_67777846 was associated with STI under WS1 of 2016 and 2017 and WS2 of 2017 using both GLM+Q and MLM. SNPs S3_13763609 and S1_74186408 were associated with RPW in WS1 and WS2 of 2017 using both GLM+Q and MLM. The identified pleiotropic lead SNPs showed significant allelic effect and significantly (*p* < 10^-8^) explained 11–25% of STI for grain weight across water deficit treatments (Table S2). Of the 16 putative pleiotropic associations, 6 associations (S4_67777846, S2_18195896, S9_57781496, S10_1402513, S6_55048997) overlapped with associations identified for the STI BLUPs across water-stressed environments (Fig. 4; File S3).

**Fig. 4.**
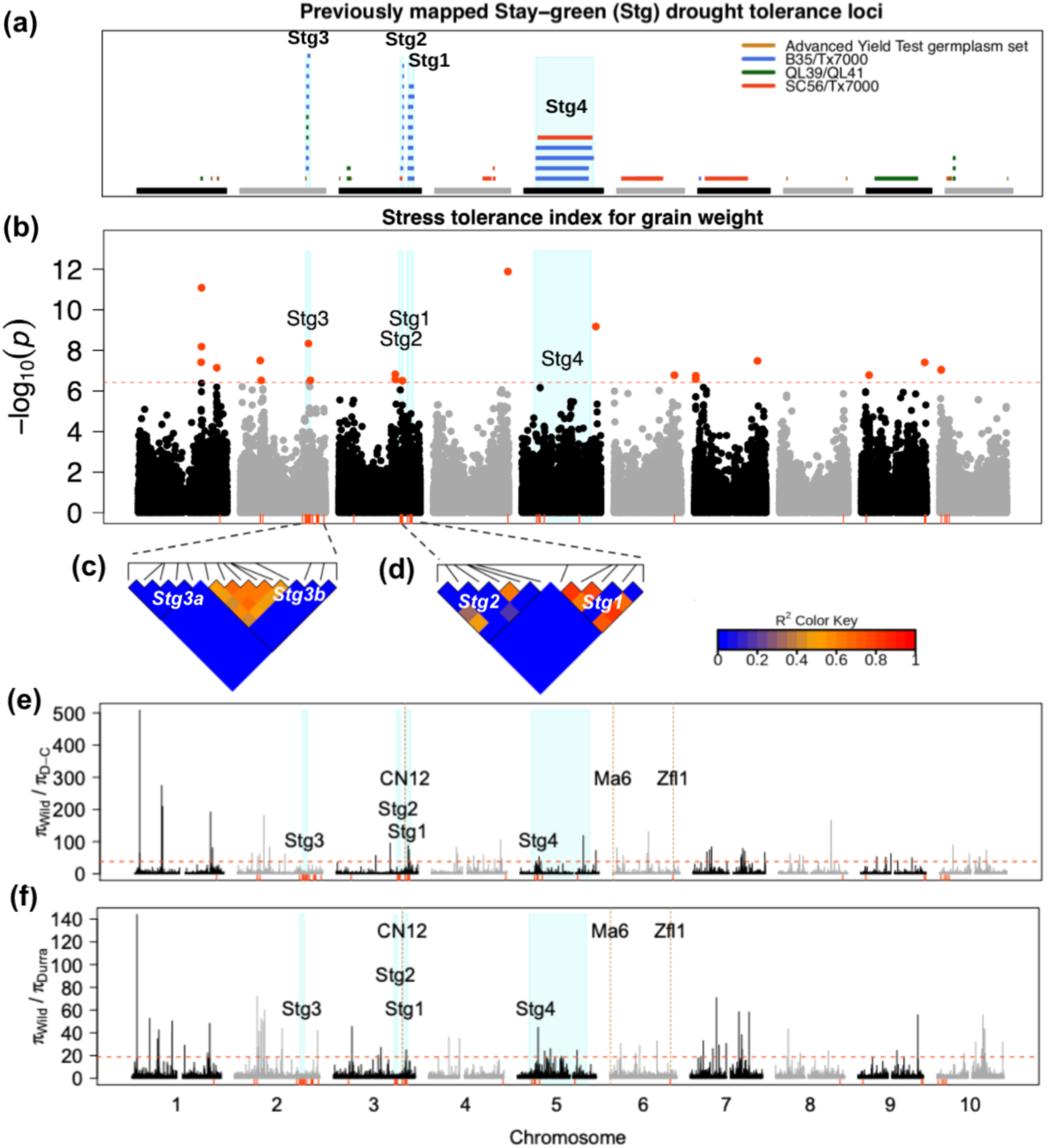
Genome-wide associations for drought tolerance and genome scans for adaptation. (a) Genomic location of the different stay-green quantitative trait loci, including *Stg1–4*, obtained from the Sorghum QTL Atlas. Light blue bars indicate the genomic position of *Stg1–4* intervals. (b) Manhattan plots of BLUPs of stress tolerance index (STI) for grain weight across pre-flowering (WS1) and post-flowering (WS2) water-stressed treatments of 2015–2017 based on GLM+Q. The horizontal red dashed line represents the Bonferroni significance threshold at 0.05, and red dots indicate lead SNPs above the threshold. Some lead SNPs colocalize within *Stg1, Stg2, Stg3,* and *Stg4* loci that are represented by light blue barplots. Rug-plots indicate the genomic position of the putative pleiotropic lead SNPs and lead SNPs at *Stg1–4* loci, significantly associated with grain weight STI and drought response variables, reduction of PW (RPW), DBM (RDBM), GrN (RGrN), PH (RPH), TGrW (RTGrW). (c) LD heatmap for lead SNPs at *Stg3a* (left triangle) and *Stg3b* (right triangle). (d) Linkage disequilibrium (LD) heatmap for lead SNPs at *Stg2* (left triangle) and *Stg1* (right triangle). (e, f) Reduction of nucleotide diversity, based on 100 kb sliding windows, (e) in durra-caudatum (D-C) and (f) in durra landraces relative to wild sorghums. Red dashed horizontal lines indicate the 99 percentile threshold for signatures of selection outliers. Rug-plots in red indicate lead SNPs for putative pleiotropic lead SNPs and lead SNPs within *Stg1–4* loci associated with drought response variables. Light blue bars indicate the genomic position of *Stg1–4* intervals.

### Drought response associations colocalizing with stay-green loci

To test the hypothesis that *Stg* loci identified from Ethiopia, we characterized the colocalization of GWAS peak SNPs with previously defined *Stg* QTL intervals as summarized in the Sorghum QTL Atlas. The interval of *Stg3* (*Stg3a* and *Stg3b*) was defined based on the introgressed region by the ICRISAT breeding program (Vadez *et al*., 2013). Of the total lead SNPs associated with STI for grain weight and drought response variables, 78 overlapped with 54 QTLs of the published *Stg* QTLs (File S4, Fig.4a,b), which represents a significant enrichment (*p* < 10^-16^). Thirty lead SNPs colocalized with known *Stg1–4* loci (Table S3). The lead SNPs colocalizing with each locus could explain up to 16% (*p* < 10^-10^, *Stg1*), 20% (*p* < 10^-13^, *Stg2*), 19% (*p* < 10^-13^, *Stg3a*), 27% (*p* < 10^-16^, *Stg3b*) and 21% (*p* < 10^-15^, *Stg4*) of the phenotypic variation across WS1 and WS2 over years based on STI BLUPs. At *Stg2*, SNP S3_56094063 was the top association (*p-*GLM < 10^-19^, *p-*MLM <10^-13^) for STI in WS2 and WS1. At *Stg3b*, S2_62095163 was the top association (*p*-GLM <10^-18^, *p*-MLM <10^-13^) with high effect for STI in WS2. The remaining lead SNP associations did not colocalize with *Stg* loci.

### Putative pleiotropic drought response associations colocalizing with stay-green loci

At each of the *Stg1–4* loci there were several associations observed across two or more drought scenarios or drought response variables (Table S3). The *Stg3a* and *Stg3b* (which are next to each other) region covered associations for STI in WS1 of 2015 and 2016, STI in WS2 of 2016 and 2017, RPW in WS1 of 2017, and RDBM in WS1 of 2016. There was a strong LD among the lead SNPs within *Stg3b* but no LD among lead SNPs within *Stg3a* (Fig. 4c). The SNP S2_62095163 was in strong LD with other lead SNPs in *Stg3b* but not in LD with lead SNPs in *Stg3a*. *Stg2* colocalized with putative pleiotropic associations for STI in WS1 of 2015 and 2017, WS2 of 2017, RGrN in WS1 of 2017, and RDBM in WS1 of 2016. The *Stg1* locus covered associations for RPW in WS1 and WS2 of 2017 and associations for STI in WS1 of 2017. At both *Stg1* and *Stg2* there was strong LD among several lead SNPs (Fig. 4d). At the *Stg4* locus there were associations for RPW in WS1 of 2017 and for STI in WS1 of 2015 and in WS2 of 2017, and moderate LD among lead SNPs (Fig. S4f).

### Evolutionary signals around drought tolerance loci

To investigate the possibility of positive selection for drought tolerance at *Stg* loci, we conducted a genome scan of pairwise nucleotide diversity (*π*) ratios for West African sorghums relative to wild relatives (i.e. outliers with high π_sorghum_/π_wild_ ratio), considering 95th and 99th percentile outliers (Fig. 4e,f; Fig. S5). Twelve of the lead SNPs associated with RPW and grain weight STI overlapped with *π* ratio outliers in durra-caudatums and durras, but not in guineas (Table 2). Colocalizations of *π* ratio outliers with *Stg1–4* loci were significantly enriched (*p* < 10^-16^). In durra-caudatums and durras, but not in guineas, some 99th percentile *π* ratio outliers were localized within *Stg1* (Fig. 4e,f; Fig. S5). We characterized the geographic distribution of two selected lead associations within each *Stg* locus to determine whether the *Stg* alleles are rare and only involved in local adaptation or are broadly involved in adaptation across sorghum landraces, beyond known sources in Ethiopia sorghums (Fig. 5a,b). The rare alleles associated with increased STI at a few selected lead SNPs within *Stg1-3* were broadly geographically distributed in sorghum landraces (Fig. 5c-h). (*Stg4* was excluded due to its large interval). However, the increased STI-associated allele at lead SNPs that overlapped with strong selection outliers were found mostly in WA sorghums (Fig. 5h), except for S3_66366589 (Fig. 5g).

**Table 2.** Pairwise-wide nucleotide diversity (π) ratio outliers overlapping with some lead SNP associations and their allele frequency in each botanical type.

| Chr | Locus | Lead SNP | Nucleotide diversity ratio |  |  | Common allele frequency |  |  |
| --- | --- | --- | --- | --- | --- | --- | --- | --- |
| | | | $\pi W/\pi DC$ | $\pi W/\pi D$ | $\pi W/\pi G$ | DC | Durra | Guinea |
| 1 | – | S1_74186408 | **18.2** | 3 | – | 0.5 | 0.5 | 0.46 |
| 2 | – | S2_18195896 | 2.4 | **9** | – | 0.49 | 0.5 | 0.47 |
| 2 | – | S2_20558788 | **61.3** | **72.2** | **54.6** | 0.49 | 0.5 | 0.45 |
| 2 | *Stg3b* | S2_71386056 | 2.1 | **7.8** | – | 0.5 | 0.5 | 0.44 |
| 2 | – | S2_76213690 | **13.4** | 3.3 | – | 0.45 | 0.26 | – |
| 3 | *Stg1* | S3_62836558 | 2.8 | 2.2 | **7.5** | 0.5 | 0.5 | 0.45 |
| 3 | *Stg1* | S3_65137990 | **15.1** | 1.7 | – | 0.5 | 0.5 | 0.46 |
| 3 | *Stg1* | S3_65430305 | **88.2** | **15.5** | **8.4** | 0.5 | 0.5 | 0.46 |
| 3 | *Stg1* | S3_66366589 | **74.6** | **25** | – | 0.5 | 0.46 | 0.46 |
| 5 | *Stg4* | S5_15215761 | **32** | 6.8 | **10.1** | 0.5 | 0.48 | 0.48 |
| 5 | *Stg4* | S5_16480120 | **27** | **13** | – | 0.45 | 0.41 | 0.48 |
| 5 | *Stg4* | S5_20251208 | 1 | 1.5 | **8.7** | 0.45 | 0.46 | 0.5 |

Bold numbers are among the 95 percentile of $\pi$ ratios.
Chr, chromosome; DC, durra-caudatum landraces; D, durra landraces; G, guinea landraces; W, wild sorghums.

**Fig. 5.**
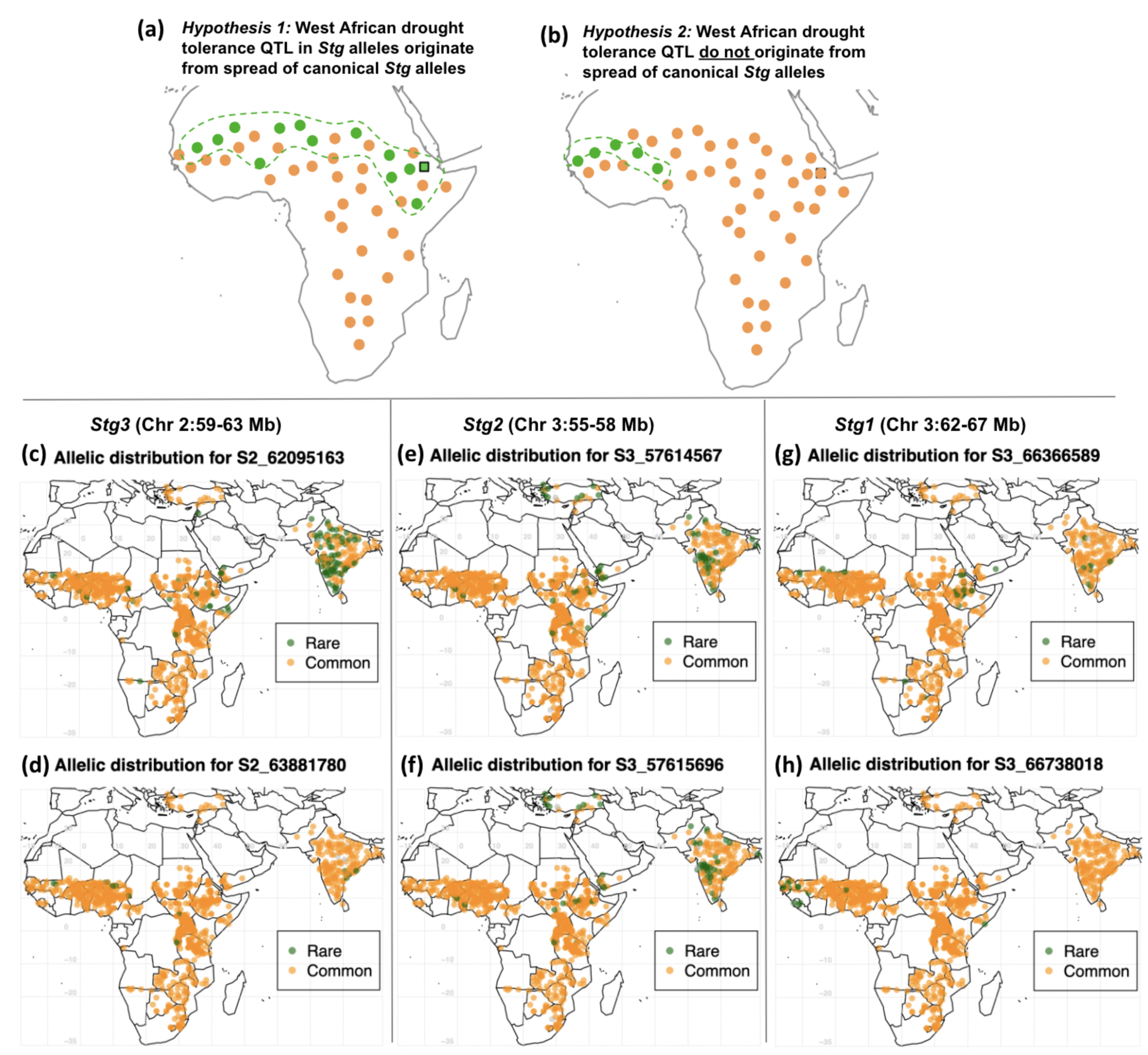
Evidence for a broad role of canonical staygreen alleles in drought adaptation. (a, b) Competing hypotheses on the origin of West African drought tolerance MTA and relationship with the canonical *Stg* alleles (titles), and graphical predictions under each hypothesis (diagrams). Under hypothesis 1 (panel a) some West African drought tolerance MTA represented *Stg* alleles that have diffused from Eastern Africa, while under hypothesis 2 (panel b) these MTA are unrelated to *Stg* alleles. The location of the known *Stg* allele source, accession IS12555 from Ethiopia, is noted by the black square. (c-h) Observed global allelic distributions at some West African drought tolerance MTA that colocalize with *Stg1–3*. (c, d) Geographic distribution of the common allele (orange) and rare allele (green) associated with increased drought tolerance of GWAS lead SNPs, S2_62095163 (c) and S2_63881780 (d) in the *Stg3* locus. (e, f) Geographic distribution of the common allele (orange) and rare allele (green) associated with increased drought tolerance of GWAS lead SNPs, S3_57614567 (e) and S3_57615696 (f) in the *Stg2* locus. (g, h) Geographic distribution of the common allele (orange) and rare allele (green) associated with increased drought tolerance of GWAS lead SNPs, S3_66366589 (g) and S3_66738018 (h) in the *Stg1* locus, respectively. The different GWAS lead SNPs above were selected based on their association with drought tolerance, proportion of variance explained, colocalization within *Stg1-3* loci, linkage disequilibrium with other lead SNPs within each *Stg1-3* locus, and availability in the GBS data for global sorghum landraces. Note, lead SNPs in *Stg4* locus were not included because of the large interval for this locus.

## DISCUSSION

### How well do managed stress trials reveal the genetics of drought tolerance in sorghum?

In this study we sought to better understand genetics of drought adaptation in sorghum, a crop that is well known for drought tolerance, but whose mechanisms of drought adaptation are not yet understood (Choudhary *et al*., 2020). We characterized a diverse panel of West African sorghum germplasm in common-garden managed drought stress field trials with the aim of balancing experimental repeatability (via the use of irrigation in off-season) with biological and breeding relevance (via the use of a field environment) (Cooper *et al*., 2014). The usefulness of managed stress experiments to understand crop evolution and improve crop resilience depends on several criteria we consider in turn:

i. *Was the intended stress applied?* Two lines of evidence support the contention that the intended drought stress was successfully imposed via irrigation management in the off-season. First, the measured soil moisture was consistently high in well-watered control treatment (FTSW ≈ 0.6), but dropped to ∼0.2 at the intended times in pre- or post-flowering drought stress treatments (Fig. 1f). The FTSW values achieved in WS1 and WS2 were similar to the critical values (∼0.2–0.5) for decreases in transpiration and leaf expansion in diverse sorghum lines (Choudhary *et al*., 2020), suggesting that a physiologically-relevant stress was experienced by the plants. Second, we observed a substantial (but not complete) reduction of yield components (∼50%; Fig. 2) under managed drought stress (WS1 and WS2 relative to WW), suggesting the stress was also agronomically relevant (Blum, 2010).
ii. *Was the stress comparable to previous stress experiments?* To be able to address this question, we included two international drought tolerance check lines, which are the canonical post-flowering (BTx642) and pre-flowering (Tx700) drought tolerant genotypes based on many studies in the U.S. and Australia (Tuinstra *et al*., 1996; Burke *et al*., 2013; Borrell *et al*., 2014b). Consistent with the idea that our managed drought stress was comparable to natural and managed drought stress in the U.S. and Australia, a strong cross-over genotype-environment interaction for grain yield of Tx7000 vs. BTx642 under pre- vs. post-flowering drought in the expected direction (Fig. 1g).
iii. *Was the timing and severity of stress comparable to that in the TPE?* Among the three criteria, this is the most difficult to assess. A formal envirotyping study, which quantifies the frequency of particular water deficits relative to crop phenology, would be necessary to address this question (Chenu *et al*., 2011; Cooper *et al*., 2014). One particular concern for off-season managed stress would be that differences in photoperiod regime relative to the TPE (i.e. the rainy season) could change in growth or developmental dynamics in a way that alters the drought response (Blum, 2010; Gano *et al*., 2021). However, the overall similarity of grain yield components in the rainy season (RF) and off-season experiments (Fig. S1a,e; Fig. S2c,d) suggest that the managed drought stress is broadly comparable to drought in the TPE. Ultimately, to rigorously test hypotheses on the similarity of off-season managed drought to the drought in the TPE, a comparison of phenotypes under managed stress to multi-environment field trials under natural drought stress will be necessary (Cooper *et al*., 1995).

### Evidence for a role of *SbZfl1* and *SbCN12* in flowering time variation and for *SbCN12* in drought adaptation

Flowering time is a critical component of geographic adaptation (Lasky *et al*., 2015; Castelletti *et al*., 2020) and a potential contributor to drought adaptation via early-flowering drought escape (Blum, 2010). Among the six canonical sorghum photoperiodic flowering genes (*Maturity1–Maturity6*) characterized in U.S. germplasm, (Murphy *et al*., 2011, 2014; Yang *et al*., 2014; Casto *et al*., 2019) we identified colocalization of associations only at *Ma2* (Fig. 3a). Instead, the top QTL mapped two *a priori* flowering time candidate loci, *SbZfl1* (chr6: 55.289–55.293 Mb) and *SbCN12* (chr3: 62.747–62.749 Mb) that are not known to underlie genetic variation in U.S. germplasm (Fig. 3; Fig. S3).

*SbZfl1* is the ortholog of maize *ZFL1/2* and rice *RLF*, which induce early flowering by activating vegetative-to-reproductive transition (Bomblies & Doebley, 2006; Rao *et al*., 2008). While *SbZfl1* variation has not been previously identified via linkage mapping (Mace *et al*., 2019), *SbZfl1* was identified as a top candidate in a recent GWAS of photoperiodic flowering rating in a Senegal regional panel (Faye *et al*., 2019). The MAF of the *SbZfl1* QTL was high (>0.4) in both WASAP and global georeferenced landraces (Fig. 3e), suggesting a common, moderate-effect variant exists at *SbZfl1*. Sorghum is a short day species, so under short days (e.g. the cool off-season in West Africa; Fig. 1b) it is expected to flower early, regardless of photoperiodism. Given *SbZfl1* was the top flowering time association under short days, *SbZfl1* may be a regulator of basic vegetative phase (BVP), the thermal time component of flowering regulation that acts independently of photoperiodic flowering regulation (Guitton *et al*., 2018). This hypothesis could explain the lack of a flowering time QTL at *SbZfl1* in a previous GWAS under long days (rainy season) in the WASAP (Faye *et al*., 2021)—subtle BVP variation at *SbZfl1* could have been masked by large-effect photoperiodic variants at *Ma6*, *SbCN8*, or other loci. However, this hypothesis would not explain the photoperiod flowering association at *SbZfl1* previously observed in Senegalese germplasm (Faye *et al*., 2019). Given inherent limitations of association studies (Korte & Farlow, 2013) and the complexity of photothermal flowering (Li *et al*., 2018b), linkage mapping and ecophysiological modeling will be needed to test these hypotheses on the role of *SbZfl1* in flowering time adaptation (Guitton *et al*., 2018; Li *et al*., 2018b).

*SbCN12* (also known as *SbFT8*) is a co-ortholog of the canonical florigen Arabidopsis *FT* gene and ortholog of maize *ZCN12* (Yang *et al*., 2014; Castelletti *et al*., 2020), which was identified as a likely sorghum florigen based on conserved sequence and expression dynamics (Yang *et al*., 2014; Wolabu *et al*., 2016). The current GWAS findings, along with previous finding that *SbCN12* explained up to 12% of variation in global nested association mapping population (Bouchet *et al*., 2017; Hu *et al*., 2019), provide strong support for the hypothesis that functional allelic variation exists at *SbCN12*. Given the early-flowering associated allele near *SbCN12* is globally rare (Fig. 3h), it may be a useful new allele for sorghum breeding programs targeting earlier flowering for stress escape. Molecular cloning of causative variants at *SbCN12* and *SbZfl1* could shed light on their role in flowering time evolution (Bomblies & Doebley, 2006; Castelletti *et al*., 2020) and facilitate development of molecular marker to recover locally-adaptive flowering time.

The evidence for a role of these flowering time genes in drought adaptation (e.g. via drought escape) is mixed. On one hand, *SbCN12* colocalized with a drought response association (RPW for WS1 vs. WW; S3_62836558; 64 kb away; Table S3), so could plausibly underlie some variation for pre-flowering drought response of this yield component. Also the same SNP near *SbCN12* was in a window of reduced π in guinea genotypes (Table 2), suggesting selection at this locus. (Note, this is not the same SNP as the rare flowering time associated variant S3_62811196, but a common variant 25 kb away). On the other hand, *SbZfl1* did not colocalize with the drought response QTL (STI, RPW, etc.; the nearest association with STI, S6_55048997, was at ∼240 kb away) and there was no evidence of positive selection around *SbZfl1* based on the *π* ratios (Fig. 4; Fig. S5). Given that causative variants at *SbCN12* and *SbZfl1* are not yet known, hypotheses on the role of these genes in drought adaptation remain speculative, but could be tested using near-isogenic lines (NILs).

### Insights on the genetics of drought adaptation in sorghum

The botanical types of sorghum vary strikingly in their morphology and geographic distribution, and based on a phytogeographic adaptation model (Vavilov, 2009). It has long been hypothesized that they vary in their drought adaptedness (Harlan & De Wet, 1972; Lasky *et al*., 2015; Wang *et al*., 2020). For instance, durra sorghums, which predominate in arid regions, are thought to be the most drought tolerant (Harlan & De Wet, 1972), while guinea sorghums, which predominate in humid regions are thought to be adapted to high humidity (De Wet *et al*., 1972). However, previous studies of large sorghum diversity panels under managed drought stress have not directly tested this hypothesis, for instance, by comparison of drought response for yield among botanical types (Vadez *et al*., 2011; Lasky *et al*., 2015; Upadhyaya *et al*., 2017). Surprisingly, in this study we find no evidence of overall differences in drought tolerance among botanical types based on the drought response of yield components (Fig. 2). These findings could be explained by one of two competing hypotheses. First, it is possible that the drought scenarios we applied do not correspond to the drought scenarios in the TPE, such that true differences in drought tolerance among botanical types were not reflected in the phenotypes. Alternatively, it may be that the major botanical types in West Africa all harbor substantial drought tolerance, presumably because droughts are common even in the higher precipitation portions of the sorghum range (Traore *et al*., 2014). In either case, our findings suggest that long-held views on differential drought adaptation among botanical types in sorghum require more formal testing.

Theoretical considerations on water use tradeoffs (Tardieu, 2012) and the apparent lack of sorghum genotypes harboring both pre- and post-flowering drought tolerance (Burke *et al*., 2013) suggest that a tradeoff might exist between early versus late stage tolerance mechanisms. However, the moderate positive correlation of the grain yield estimates under pre- and post-flowering drought (e.g. for GrW or STI; Fig. S2a) suggest no major physiological tradeoff for tolerance to these contrasting drought scenarios, at least at this broad scale of diversity. Colocalization of MTA for drought tolerance related traits in WS1 and WS2 would provide further evidence for genetic factors that contribute pleiotropically to both pre- and post-flowering drought tolerance. Consistent with this hypothesis, sixteen distinct MTAs (Table S2) were detected for drought-related traits (mostly STI) under both the pre- and post-flowering drought treatments, including one (S3_56094063) that colocalized with *Stg1–4* loci (Table S3).

Among the positive pleiotropic associations, the STI MTA at S4_67777846 may be the most interesting candidate for further study, given that it had the highest PVE estimate (25%) in both pre- and post-flowering water stress over two years. This MTA did not colocalize with *Stg* QTLs or other *a priori* candidate genes, and there were no obvious *post hoc* candidate genes near the SNP, so we have no hypothesis on the biological basis of this association at this point. If confirmed, positive pleiotropic drought tolerance QTLs, which could contribute to yield stability across drought scenarios, would be of great interest for breeding of broadly-adapted climate-resilient varieties and help elucidate mechanisms that circumvent potential tradeoffs (Tardieu, 2012).

Another question that motivated our study was whether canonical *Stg* alleles, which were originally discovered in Ethiopia-derived materials (BTx642) (Borrell *et al*., 2014a), are also present in West African landraces (Fig. 5a). The hypothesis that canonical *Stg* alleles have a broad role in drought adaptation across Africa is plausible since it is well established that durra sorghums diffused from Ethiopia across the Sahel to West Africa (Harlan & Stemler, 1976; Morris *et al*., 2013). Indeed, we observe a statistically significant enrichment of drought tolerance related MTA colocalized with canonical *Stg* QTL intervals, which provides preliminary support for the shared *Stg* hypothesis (File S4; Table S3). Most notable among these are the highly significant association for grain weight STI under post-flowering drought at *Stg3* (S2_62095163) and *Stg2* (S3_57614567). Further, the West Africa drought tolerance associated alleles in the *Stg* intervals are found in Ethiopia, as would be expected if they were shared across Africa. While these findings are suggestive, they are not sufficient to exclude the alternate hypothesis (Fig. 5b) that West African drought tolerance loci are unrelated to Ethiopian-origin *Stg* alleles. Testing this hypothesis conclusively would require positional cloning of the West Africa drought tolerance QTL and *Stg* alleles.

The final hypothesis we considered was that drought tolerance alleles underlie drought adaptation *per se* and were subject to positive selection. This finding was supported by significant enrichment for colocalization of selection outliers with *Stg* QTLs and common allele frequencies of lead SNPs overlapping with selection outliers in durra-caudatums and durras relative to guineas (Fig. 4e,f; Fig. S5; Table 2). As with the other findings, the development of NILs and the validation of major effect QTL in breeding populations (Borrell *et al*., 2014b,a) will be crucial to rigorously test the proposed role of QTL in these genomic regions for drought adaptation. Overall, our findings support the long-standing hypothesis that genetic variation for drought tolerance exists in West African sorghum, and provide preliminary evidence for a broad role of canonical *Stg* drought tolerance alleles across Africa.

## ACKNOWLEDGMENTS

This study is made possible by the support of the American People provided to the Feed the Future Innovation Lab for Collaborative Research on Sorghum and Millet through the United States Agency for International Development (USAID) under Cooperative Agreement No. AID- OAA-A-13-00047. The contents are the sole responsibility of the authors and do not necessarily reflect the views of USAID or the United States Government. Analyses were made possible by Beocat High-Performance Computing Cluster at Kansas State University.

## Author Contributions

GPM, DF, NC, BS: design of the research; EAA, BS, CD: performance of the research; JMF, EAA, BS, GPM: data analysis, collection, or interpretation; JMF, GPM: writing the manuscript.

## SUPPORTING INFORMATION

**Fig. S1.**
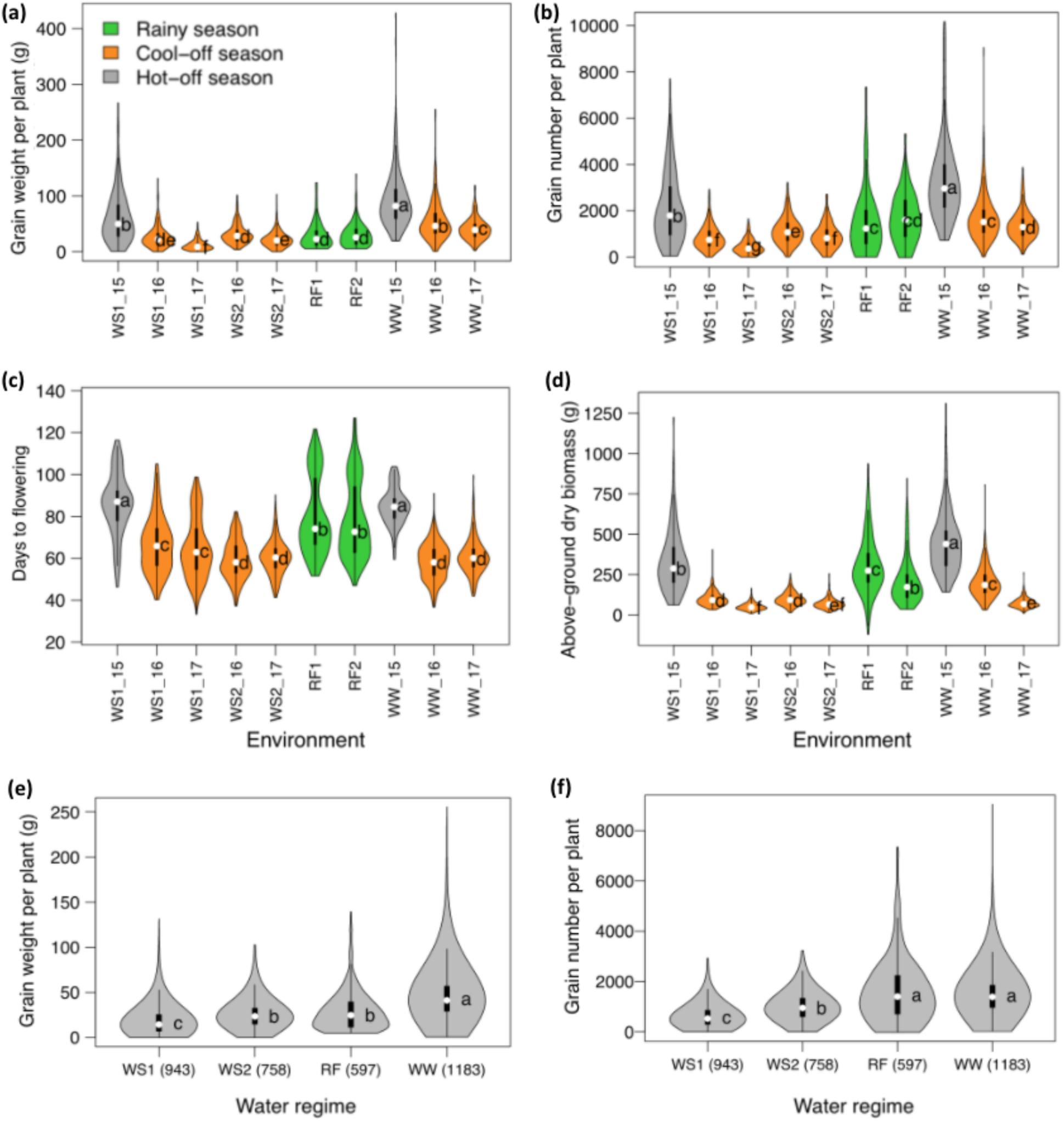
Summary of phenotypic data across environments. (a-d) Effect of water deficit on grain weight per plant (a), grain number per plant (b), days to flowering (c), and above-ground dry biomass (d) of accessions in each environment across the different water regimes (WS1, pre-flowering water stress; WS2, post-flowering water stress; RF, rainfed; WW, well-watered). The environments under rainfed (RF1 and RF2), cool off-season, and hot off-season conditions are colored in green, orange, and gray, respectively. (e, f) Average values for grain weight per plant (e) and grain number per plant (f) in each water regime, excluding the 2015 data. Letters within violin plots indicate the TukeyHSD statistical difference at α = 0.05.

**Fig. S2.**
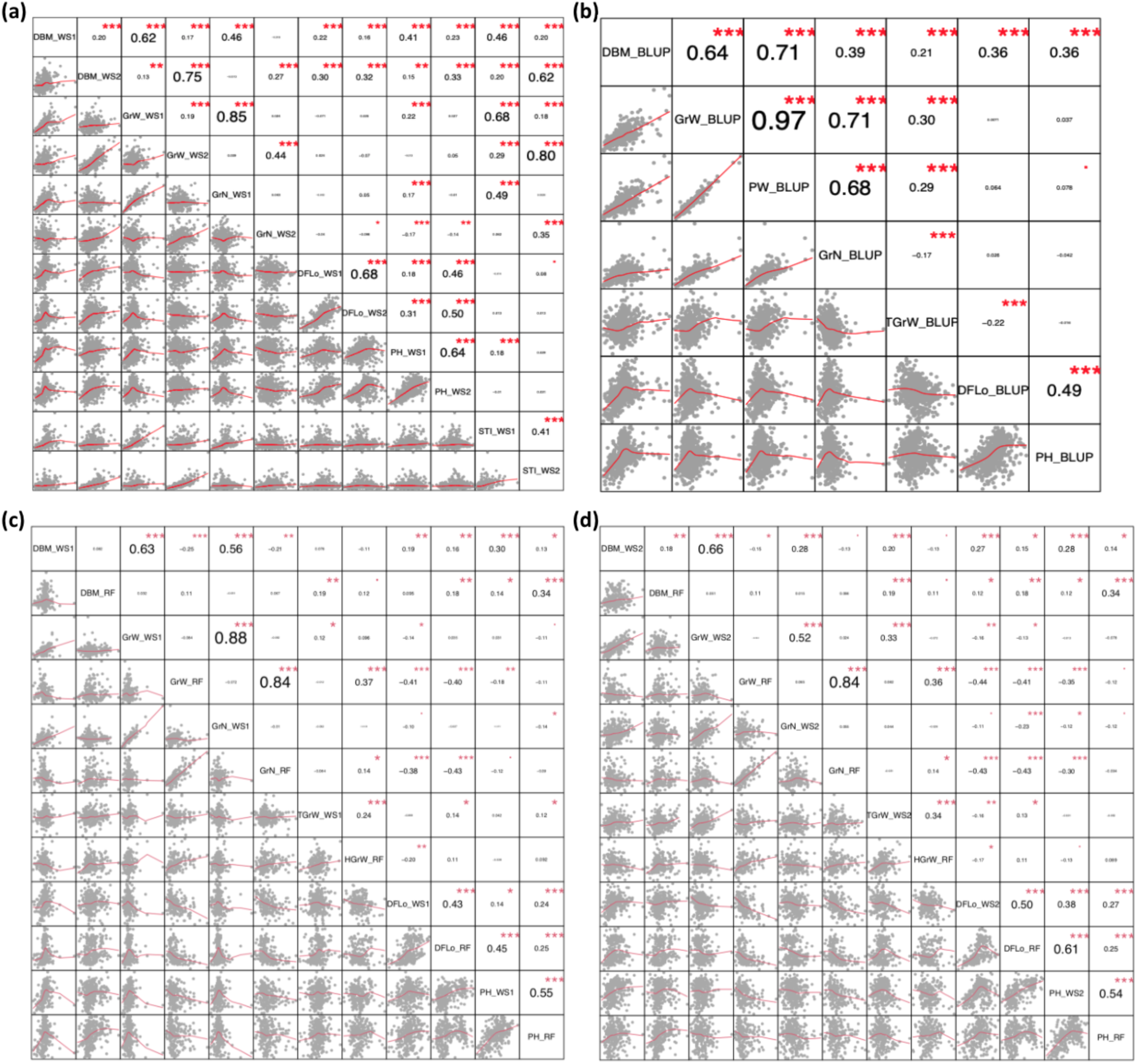
Relationship among phenotypes across environments. (a) Correlations for yield components based on BLUP values in pre-flowering (WS1) and BLUP values in post-flowering (WS2) water stress environments. (b) Correlations for yield components based on BLUP values across all environments. (c, d) Correlations for yield components based on BLUP values in rainfed (RF) environments relative to WS1 (c) and WS2 (d) water stress environments. DBM, above-ground dry biomass; GrW, grain weight per plant; PW, panicle weight per plant; GrN, grain number per plant; TGrW, thousand-grain weight; DFLo, days to flowering; and PH, plant height; HGrW, hundred grain weight.

**Fig. S3.**
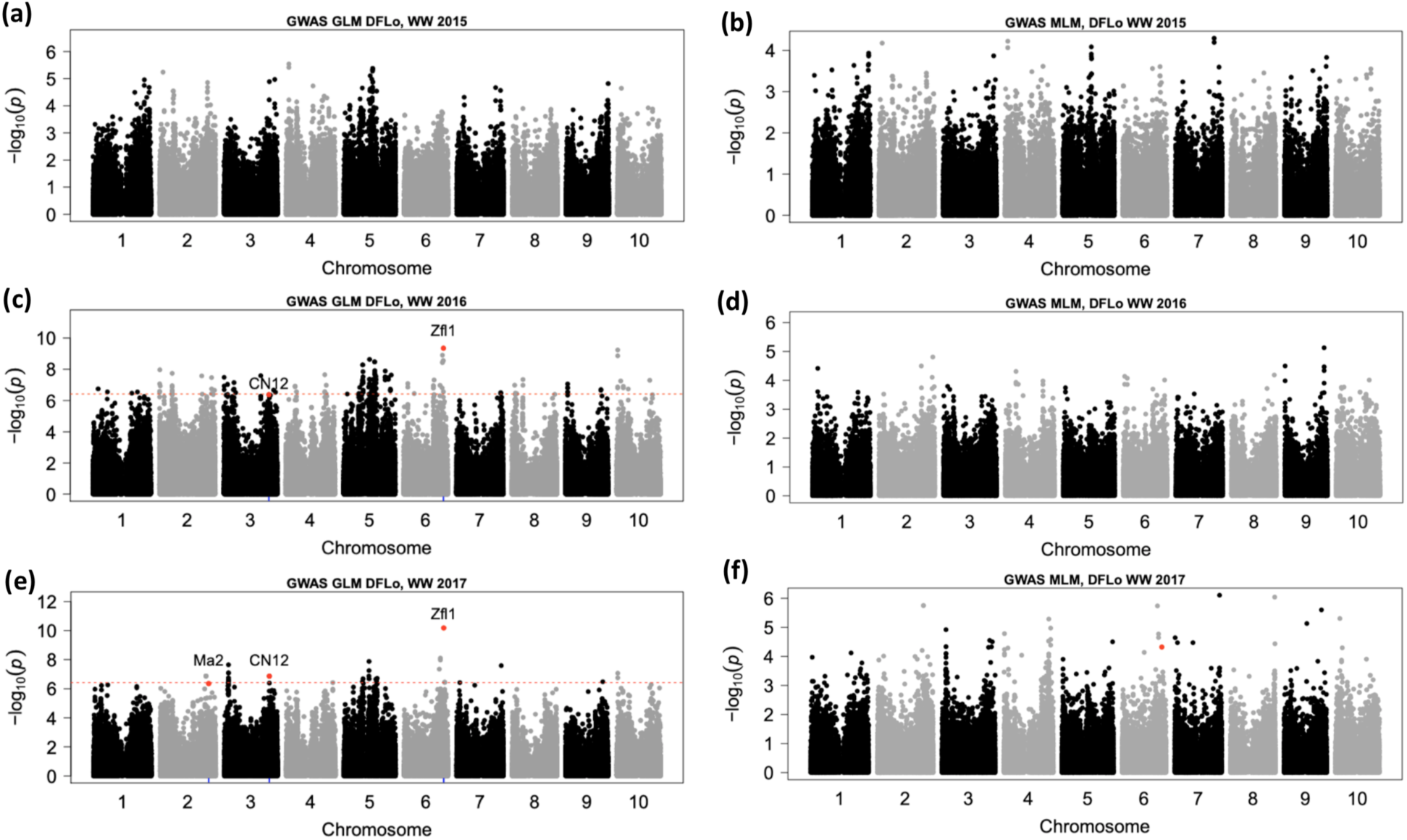
Genome-wide associations for days to flowering (DFLo) under well-watered environments over three years. (a, b) Manhattan plots for days to flowering in 2015 using (a) general linear model with principal components (GLM+Q) and (b) mixed-linear model (MLM). (c, d) Manhattan plots for days to flowering in 2016 using (c) GLM+Q and (d) MLM. (e, f) Manhattan plots for days to flowering in 2017 using (e) GLM+Q and (f) MLM. Horizontal dashed line indicates the Bonferroni correction at 0.05. Red dots indicate peak SNPs colocalizing (based on 150 kb cutoff) with flowering time candidate genes.

**Fig. S4.**
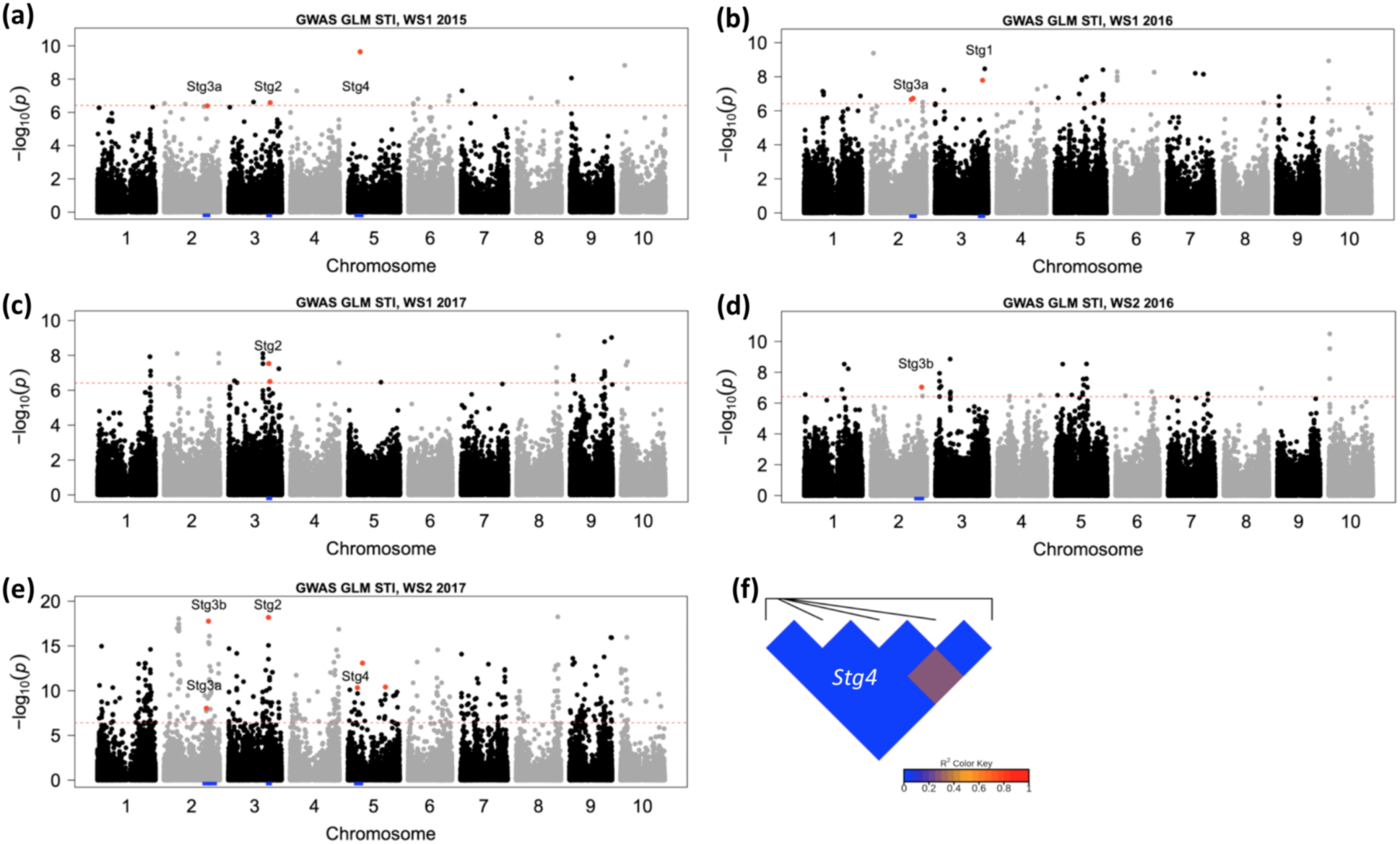
Genome-wide associations for stress tolerance index (STI) for grain weight under drought stress conditions. (a-e) Manhattan plots of STI in pre-flowering drought (WS1) of (a) 2015, (b) 2016, and (c) 2017 and post-flowering drought (WS2) of (d) 2016 and (e) 2017, based on the general linear model with principal components covariates (GLM+Q). The horizontal red dashed line represents the Bonferroni significance threshold at 0.05. Red dots indicate lead SNPs colocalizing within *Stg1–4* loci, which are represented by blue rug plots. (f) Heatmap for lead SNPs, S5_13190947, S5_15215761, S5_15916423, S5_16480120, S5_20251208, S5_52255304 at *Stg4*, associated with STI and drought response for panicle weight (RPW) and represented on Table S3.

**Fig. S5.**
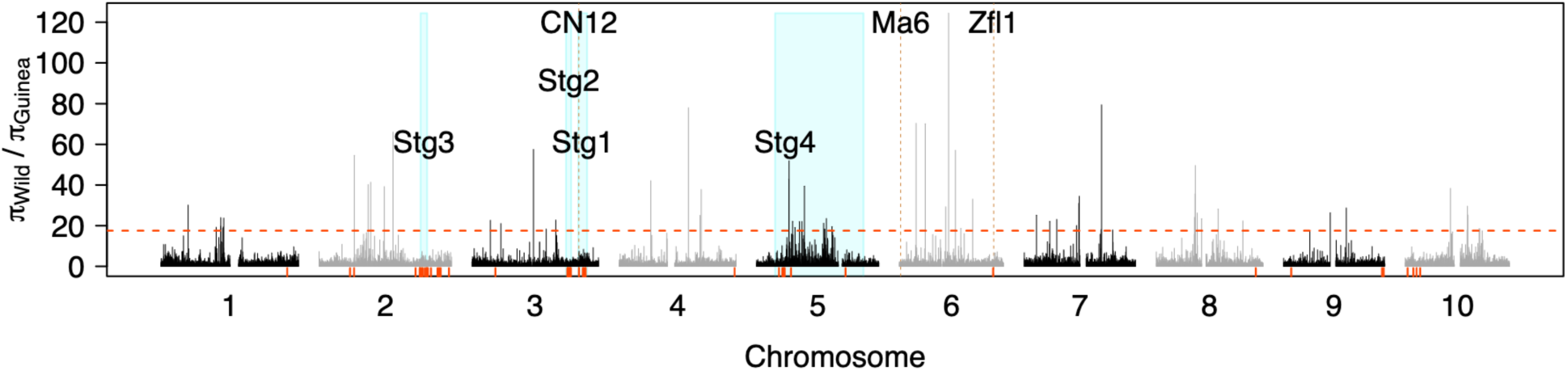
Reduction of nucleotide diversity in guinea landraces relative to wild sorghums, based on 100 kb sliding windows. Red dashed horizontal lines indicate the 99 percentile threshold for signatures of selection outliers. Barplots in skyblue indicate the genomic position of the stay-green *Stg1–4* loci. Rug-plots in red indicate lead SNPs for putative pleiotropic QTL and lead SNPs within *Stg1–4* loci associated with drought response variables.

**Table S1:**
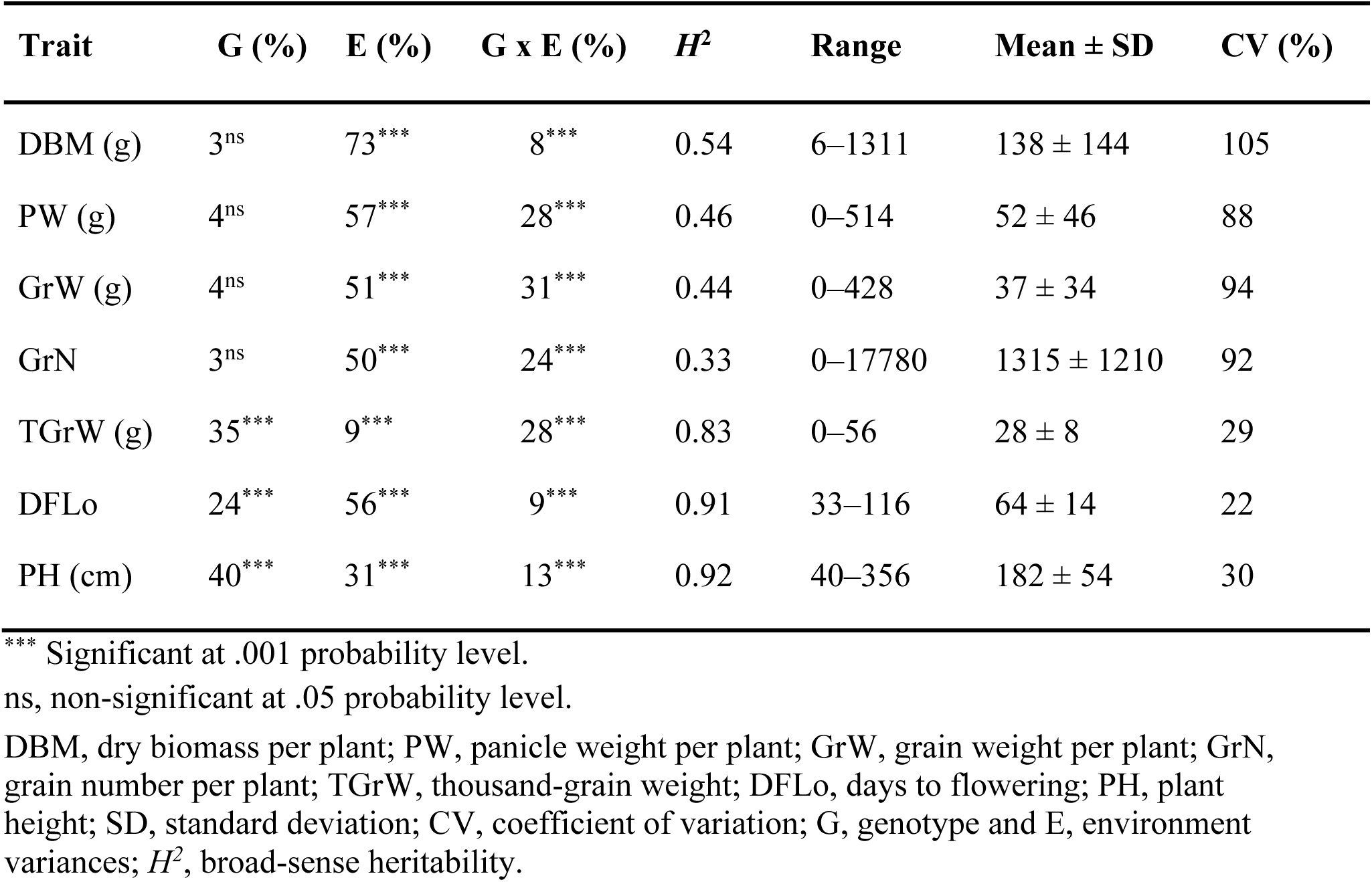
Descriptive statistics, variance components, and broad-sense heritability (*H*^2^) of yield components across all managed water stress environments.

| Trait | G (%) | E (%) | G x E (%) | $H^2$ | Range | Mean ± SD | CV (%) |
| --- | --- | --- | --- | --- | --- | --- | --- |
| DBM (g) | 3<sup>ns</sup> | 73<sup>***</sup> | 8<sup>***</sup> | 0.54 | 6–1311 | 138 ± 144 | 105 |
| PW (g) | 4<sup>ns</sup> | 57<sup>***</sup> | 28<sup>***</sup> | 0.46 | 0–514 | 52 ± 46 | 88 |
| GrW (g) | 4<sup>ns</sup> | 51<sup>***</sup> | 31<sup>***</sup> | 0.44 | 0–428 | 37 ± 34 | 94 |
| GrN | 3<sup>ns</sup> | 50<sup>***</sup> | 24<sup>***</sup> | 0.33 | 0–17780 | 1315 ± 1210 | 92 |
| TGrW (g) | 35<sup>***</sup> | 9<sup>***</sup> | 28<sup>***</sup> | 0.83 | 0–56 | 28 ± 8 | 29 |
| DFLo | 24<sup>***</sup> | 56<sup>***</sup> | 9<sup>***</sup> | 0.91 | 33–116 | 64 ± 14 | 22 |
| PH (cm) | 40<sup>***</sup> | 31<sup>***</sup> | 13<sup>***</sup> | 0.92 | 40–356 | 182 ± 54 | 30 |

\*\*\* Significant at .001 probability level.
ns, non-significant at .05 probability level.
DBM, dry biomass per plant; PW, panicle weight per plant; GrW, grain weight per plant; GrN, grain number per plant; TGrW, thousand-grain weight; DFLo, days to flowering; PH, plant height; SD, standard deviation; CV, coefficient of variation; G, genotype and E, environment variances; $H^2$ , broad-sense heritability.

**Table S2:**
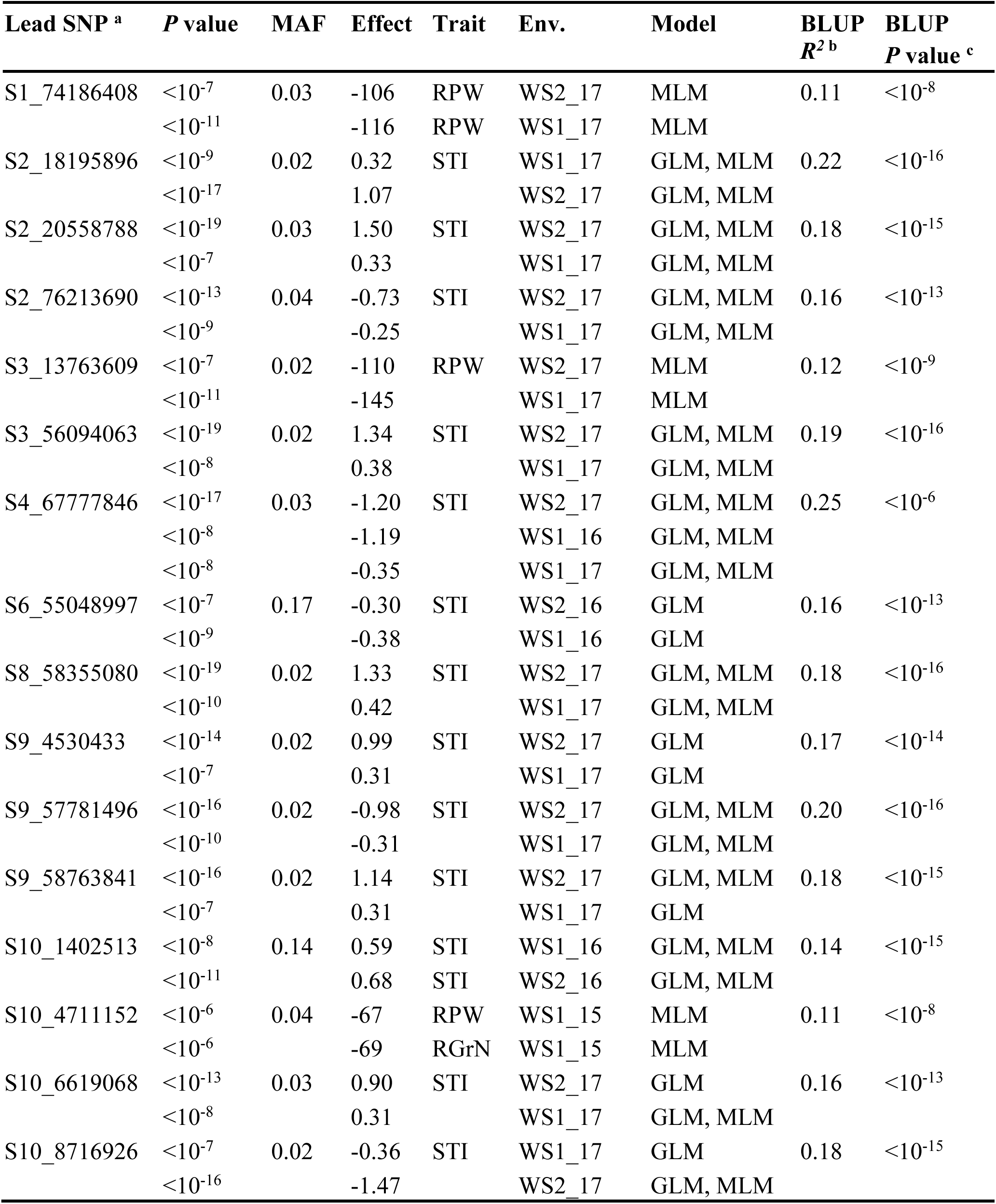
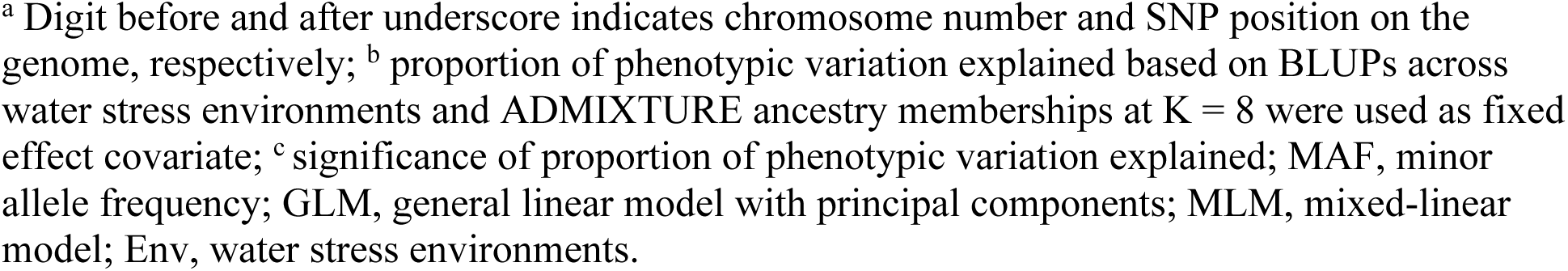
Putative pleiotropic lead SNP associations with reduction of yield components and stress tolerance index (STI) for grain weight in separate and across water stress environments.

| Lead SNP <sup>a</sup> | *P* value | MAF | Effect | Trait | Env. | Model | BLUP<br>*R*<sup>2</sup> <sup>b</sup> | BLUP<br>*P* value <sup>c</sup> |
| --- | --- | --- | --- | --- | --- | --- | --- | --- |
| S1_74186408 | <10<sup>-7</sup> | 0.03 | -106 | RPW | WS2_17 | MLM | 0.11 | <10<sup>-8</sup> |
|  | <10<sup>-11</sup> |  | -116 | RPW | WS1_17 | MLM |  |  |
| S2_18195896 | <10<sup>-9</sup> | 0.02 | 0.32 | STI | WS1_17 | GLM, MLM | 0.22 | <10<sup>-16</sup> |
|  | <10<sup>-17</sup> |  | 1.07 |  | WS2_17 | GLM, MLM |  |  |
| S2_20558788 | <10<sup>-19</sup> | 0.03 | 1.50 | STI | WS2_17 | GLM, MLM | 0.18 | <10<sup>-15</sup> |
|  | <10<sup>-7</sup> |  | 0.33 |  | WS1_17 | GLM, MLM |  |  |
| S2_76213690 | <10<sup>-13</sup> | 0.04 | -0.73 | STI | WS2_17 | GLM, MLM | 0.16 | <10<sup>-13</sup> |
|  | <10<sup>-9</sup> |  | -0.25 |  | WS1_17 | GLM, MLM |  |  |
| S3_13763609 | <10<sup>-7</sup> | 0.02 | -110 | RPW | WS2_17 | MLM | 0.12 | <10<sup>-9</sup> |
|  | <10<sup>-11</sup> |  | -145 |  | WS1_17 | MLM |  |  |
| S3_56094063 | <10<sup>-19</sup> | 0.02 | 1.34 | STI | WS2_17 | GLM, MLM | 0.19 | <10<sup>-16</sup> |
|  | <10<sup>-8</sup> |  | 0.38 |  | WS1_17 | GLM, MLM |  |  |
| S4_67777846 | <10<sup>-17</sup> | 0.03 | -1.20 | STI | WS2_17 | GLM, MLM | 0.25 | <10<sup>-6</sup> |
|  | <10<sup>-8</sup> |  | -1.19 |  | WS1_16 | GLM, MLM |  |  |
|  | <10<sup>-8</sup> |  | -0.35 |  | WS1_17 | GLM, MLM |  |  |
| S6_55048997 | <10<sup>-7</sup> | 0.17 | -0.30 | STI | WS2_16 | GLM | 0.16 | <10<sup>-13</sup> |
|  | <10<sup>-9</sup> |  | -0.38 |  | WS1_16 | GLM |  |  |
| S8_58355080 | <10<sup>-19</sup> | 0.02 | 1.33 | STI | WS2_17 | GLM, MLM | 0.18 | <10<sup>-16</sup> |
|  | <10<sup>-10</sup> |  | 0.42 |  | WS1_17 | GLM, MLM |  |  |
| S9_4530433 | <10<sup>-14</sup> | 0.02 | 0.99 | STI | WS2_17 | GLM | 0.17 | <10<sup>-14</sup> |
|  | <10<sup>-7</sup> |  | 0.31 |  | WS1_17 | GLM |  |  |
| S9_57781496 | <10<sup>-16</sup> | 0.02 | -0.98 | STI | WS2_17 | GLM, MLM | 0.20 | <10<sup>-16</sup> |
|  | <10<sup>-10</sup> |  | -0.31 |  | WS1_17 | GLM, MLM |  |  |
| S9_58763841 | <10<sup>-16</sup> | 0.02 | 1.14 | STI | WS2_17 | GLM, MLM | 0.18 | <10<sup>-15</sup> |
|  | <10<sup>-7</sup> |  | 0.31 |  | WS1_17 | GLM |  |  |
| S10_1402513 | <10<sup>-8</sup> | 0.14 | 0.59 | STI | WS1_16 | GLM, MLM | 0.14 | <10<sup>-15</sup> |
|  | <10<sup>-11</sup> |  | 0.68 | STI | WS2_16 | GLM, MLM |  |  |
| S10_4711152 | <10<sup>-6</sup> | 0.04 | -67 | RPW | WS1_15 | MLM | 0.11 | <10<sup>-8</sup> |
|  | <10<sup>-6</sup> |  | -69 | RGrN | WS1_15 | MLM |  |  |
| S10_6619068 | <10<sup>-13</sup> | 0.03 | 0.90 | STI | WS2_17 | GLM | 0.16 | <10<sup>-13</sup> |
|  | <10<sup>-8</sup> |  | 0.31 |  | WS1_17 | GLM, MLM |  |  |
| S10_8716926 | <10<sup>-7</sup> | 0.02 | -0.36 | STI | WS1_17 | GLM | 0.18 | <10<sup>-15</sup> |
|  | <10<sup>-16</sup> |  | -1.47 |  | WS2_17 | GLM, MLM |  |  |

<sup>a</sup> Digit before and after underscore indicates chromosome number and SNP position on the genome, respectively; <sup>b</sup> proportion of phenotypic variation explained based on BLUPs across water stress environments and ADMIXTURE ancestry memberships at K = 8 were used as fixed effect covariate; <sup>c</sup> significance of proportion of phenotypic variation explained; MAF, minor allele frequency; GLM, general linear model with principal components; MLM, mixed-linear model; Env, water stress environments.

**Table S3:**
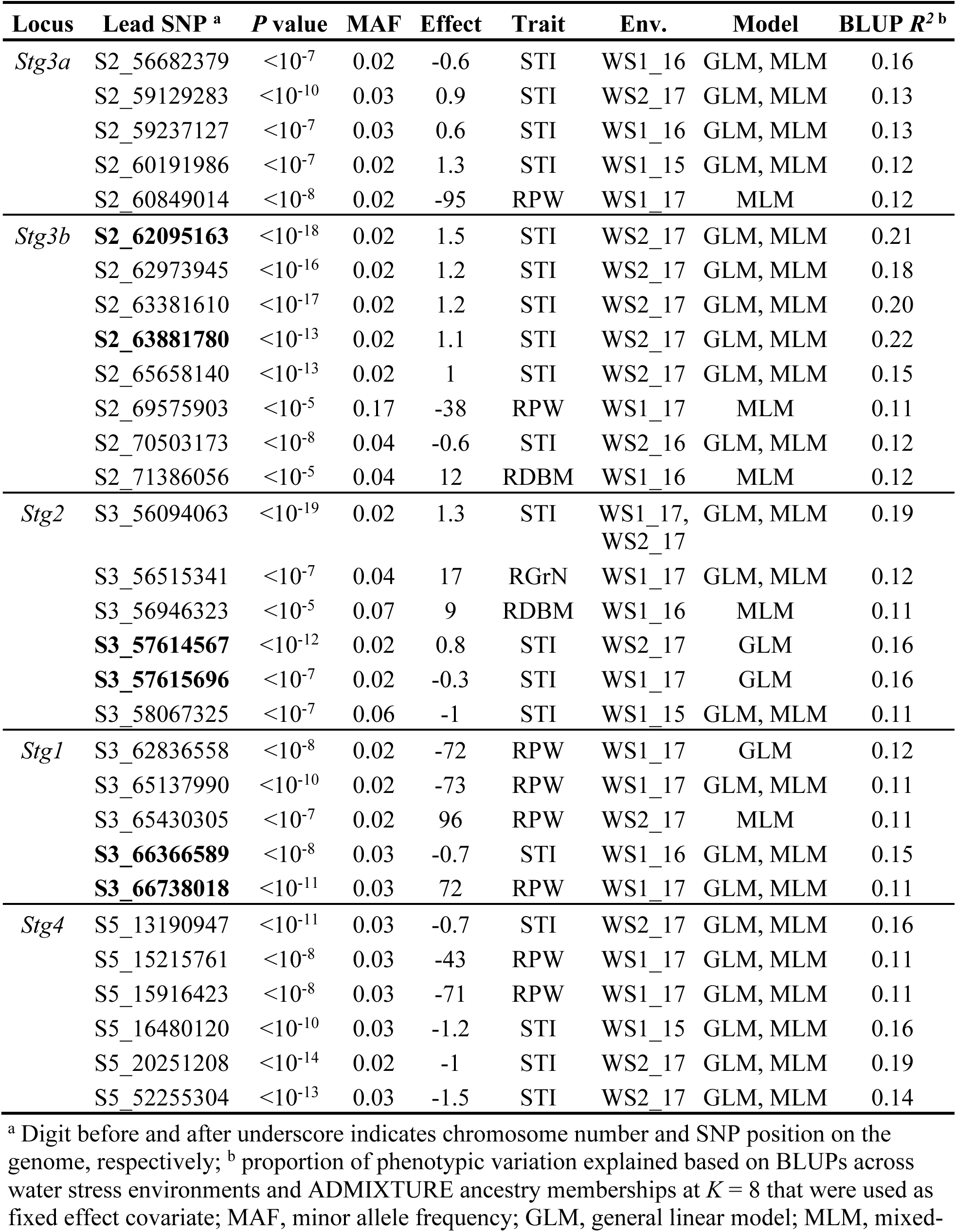
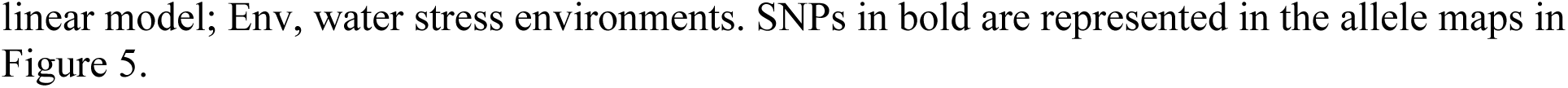
Lead SNP associations within *Stg1–4* loci for reduction of yield components and stress tolerance index for grain weight in separate and across water stress environments.

| Locus | Lead SNP <sup>a</sup> | *P* value | MAF | Effect | Trait | Env. | Model | BLUP *R*<sup>2</sup> <sup>b</sup> |
| --- | --- | --- | --- | --- | --- | --- | --- | --- |
| *Stg3a* | S2_56682379 | <10<sup>-7</sup> | 0.02 | -0.6 | STI | WS1_16 | GLM, MLM | 0.16 |
|  | S2_59129283 | <10<sup>-10</sup> | 0.03 | 0.9 | STI | WS2_17 | GLM, MLM | 0.13 |
|  | S2_59237127 | <10<sup>-7</sup> | 0.03 | 0.6 | STI | WS1_16 | GLM, MLM | 0.13 |
|  | S2_60191986 | <10<sup>-7</sup> | 0.02 | 1.3 | STI | WS1_15 | GLM, MLM | 0.12 |
|  | S2_60849014 | <10<sup>-8</sup> | 0.02 | -95 | RPW | WS1_17 | MLM | 0.12 |
| *Stg3b* | **S2_62095163** | <10<sup>-18</sup> | 0.02 | 1.5 | STI | WS2_17 | GLM, MLM | 0.21 |
|  | S2_62973945 | <10<sup>-16</sup> | 0.02 | 1.2 | STI | WS2_17 | GLM, MLM | 0.18 |
|  | S2_63381610 | <10<sup>-17</sup> | 0.02 | 1.2 | STI | WS2_17 | GLM, MLM | 0.20 |
|  | **S2_63881780** | <10<sup>-13</sup> | 0.02 | 1.1 | STI | WS2_17 | GLM, MLM | 0.22 |
|  | S2_65658140 | <10<sup>-13</sup> | 0.02 | 1 | STI | WS2_17 | GLM, MLM | 0.15 |
|  | S2_69575903 | <10<sup>-5</sup> | 0.17 | -38 | RPW | WS1_17 | MLM | 0.11 |
|  | S2_70503173 | <10<sup>-8</sup> | 0.04 | -0.6 | STI | WS2_16 | GLM, MLM | 0.12 |
|  | S2_71386056 | <10<sup>-5</sup> | 0.04 | 12 | RDBM | WS1_16 | MLM | 0.12 |
| *Stg2* | S3_56094063 | <10<sup>-19</sup> | 0.02 | 1.3 | STI | WS1_17, WS2_17 | GLM, MLM | 0.19 |
|  | S3_56515341 | <10<sup>-7</sup> | 0.04 | 17 | RGrN | WS1_17 | GLM, MLM | 0.12 |
|  | S3_56946323 | <10<sup>-5</sup> | 0.07 | 9 | RDBM | WS1_16 | MLM | 0.11 |
|  | **S3_57614567** | <10<sup>-12</sup> | 0.02 | 0.8 | STI | WS2_17 | GLM | 0.16 |
|  | **S3_57615696** | <10<sup>-7</sup> | 0.02 | -0.3 | STI | WS1_17 | GLM | 0.16 |
|  | S3_58067325 | <10<sup>-7</sup> | 0.06 | -1 | STI | WS1_15 | GLM, MLM | 0.11 |
| *Stg1* | S3_62836558 | <10<sup>-8</sup> | 0.02 | -72 | RPW | WS1_17 | GLM | 0.12 |
|  | S3_65137990 | <10<sup>-10</sup> | 0.02 | -73 | RPW | WS1_17 | GLM, MLM | 0.11 |
|  | S3_65430305 | <10<sup>-7</sup> | 0.02 | 96 | RPW | WS2_17 | MLM | 0.11 |
|  | **S3_66366589** | <10<sup>-8</sup> | 0.03 | -0.7 | STI | WS1_16 | GLM, MLM | 0.15 |
|  | **S3_66738018** | <10<sup>-11</sup> | 0.03 | 72 | RPW | WS1_17 | GLM, MLM | 0.11 |
| *Stg4* | S5_13190947 | <10<sup>-11</sup> | 0.03 | -0.7 | STI | WS2_17 | GLM, MLM | 0.16 |
|  | S5_15215761 | <10<sup>-8</sup> | 0.03 | -43 | RPW | WS1_17 | GLM, MLM | 0.11 |
|  | S5_15916423 | <10<sup>-8</sup> | 0.03 | -71 | RPW | WS1_17 | GLM, MLM | 0.11 |
|  | S5_16480120 | <10<sup>-10</sup> | 0.03 | -1.2 | STI | WS1_15 | GLM, MLM | 0.16 |
|  | S5_20251208 | <10<sup>-14</sup> | 0.02 | -1 | STI | WS2_17 | GLM, MLM | 0.19 |
|  | S5_52255304 | <10<sup>-13</sup> | 0.03 | -1.5 | STI | WS2_17 | GLM, MLM | 0.14 |

<sup>a</sup> Digit before and after underscore indicates chromosome number and SNP position on the genome, respectively; <sup>b</sup> proportion of phenotypic variation explained based on BLUPs across water stress environments and ADMIXTURE ancestry memberships at *K* = 8 that were used as fixed effect covariate; MAF, minor allele frequency; GLM, general linear model; MLM, mixed-
linear model; Env, water stress environments. SNPs in bold are represented in the allele maps in Figure 5.

## REFERENCES

Alexander DH, Novembre J, Lange K. 2009. Fast model-based estimation of ancestry in unrelated individuals. Genome Research 19: 1655–1664.

Blum A. 2010. Plant Breeding for Water-Limited Environments. New York: Springer Publishing.

Bomblies K, Doebley JF. 2006. Pleiotropic effects of the duplicate maize FLORICAULA/LEAFY genes zfl1 and zfl2 on traits under selection during maize domestication. Genetics 172: 519–531.

Borrell AK, Mullet JE, George-Jaeggli B, Oosterom EJ van, Hammer GL, Klein PE, Jordan DR. 2014a. Drought adaptation of stay-green sorghum is associated with canopy development, leaf anatomy, root growth, and water uptake. Journal of Experimental Botany 65: 6251–6263.

Borrell AK, van Oosterom EJ, Mullet JE, George-Jaeggli B, Jordan DR, Klein PE, Hammer GL. 2014b. Stay-green alleles individually enhance grain yield in sorghum under drought by modifying canopy development and water uptake patterns. New Phytologist 203: 817–830.

Bouchet S, Olatoye MO, Marla SR, Perumal R, Tesso T, Yu J, Tuinstra M, Morris GP. 2017. Increased power to dissect adaptive traits in global sorghum diversity using a nested association mapping population. Genetics 206: 573–585.

Burke JJ, Chen J, Burow G, Mechref Y, Rosenow D, Payton P, Xin Z, Hayes CM. 2013. Leaf dhurrin content is a quantitative measure of the level of pre-and post-flowering drought tolerance in sorghum. Crop Science 53: 1056–1065.

Castelletti S, Coupel-Ledru A, Granato I, Palaffre C, Cabrera-Bosquet L, Tonelli C, Nicolas SD, Tardieu F, Welcker C, Conti L. 2020. Maize adaptation across temperate climates was obtained via expression of two florigen genes. PLOS Genetics 16: e1008882.

Casto AL, Mattison AJ, Olson SN, Thakran M, Rooney WL, Mullet JE. 2019. Maturity2, a novel regulator of flowering time in Sorghum bicolor, increases expression of SbPRR37 and SbCO in long days delaying flowering. PLOS ONE 14: e0212154.

Chenu K, Cooper M, Hammer GL, Mathews KL, Dreccer MF, Chapman SC. 2011. Environment characterization as an aid to wheat improvement: interpreting genotype–environment interactions by modelling water-deficit patterns in North-Eastern Australia. Journal of Experimental Botany 62: 1743–1755.

Choudhary S, Guha A, Kholova J, Pandravada A, Messina CD, Cooper M, Vadez V. 2020. Maize, sorghum, and pearl millet have highly contrasting species strategies to adapt to water stress and climate change-like conditions. Plant Science 295: 110297.

Cooper M, Messina CD, Podlich D, Totir LR, Baumgarten A, Hausmann NJ, Wright D, Graham G. 2014. Predicting the future of plant breeding: complementing empirical evaluation with genetic prediction. Crop and Pasture Science 65: 311–336.

Cooper M, Woodruff DR, Eisemann RL, Brennan PS, DeLacy IH. 1995. A selection strategy to accommodate genotype-by-environment interaction for grain yield of wheat: managed-environments for selection among genotypes. Theoretical and Applied Genetics 90: 492–502.

Danecek P, Auton A, Abecasis G, Albers CA, Banks E, DePristo MA, Handsaker RE, Lunter G, Marth GT, Sherry ST, et al.2011. The variant call format and VCFtools. Bioinformatics 27: 2156–2158.

De Wet JMJ, Harlan JR, Kurmarohita B. 1972. Origin and evolution of guinea sorghums. East African Agricultural and Forestry Journal 38: 114–119.

Faye JM, Maina F, Akata EA, Sine B, Diatta C, Mamadou A, Marla S, Bouchet S, Teme N, Rami J-F, et al.2021. A genomics resource for genetics, physiology, and breeding of West African sorghum. The Plant Genome n/a: e20075.

Faye JM, Maina F, Hu Z, Fonceka D, Cisse N, Morris GP. 2019. Genomic signatures of adaptation to Sahelian and Soudanian climates in sorghum landraces of Senegal. Ecology and Evolution 9: 1–14.

Gano B, Dembele JSB, Tovignan TK, Sine B, Vadez V, Diouf D, Audebert A. 2021. Adaptation responses to early drought stress of West Africa sorghum varieties. Agronomy 11: 443.

Guitton B, Théra K, Tékété ML, Pot D, Kouressy M, Témé N, Rami J-F, Vaksmann M. 2018. Integrating genetic analysis and crop modeling: A major QTL can finely adjust photoperiod-sensitive sorghum flowering. Field Crops Research 221: 7–18.

Harlan JR, De Wet JJM. 1972. A simplified classification of cultivated sorghum. Crop Science 12: 172–176.

Harlan JR, Stemler A. 1976. The Races of Sorghum in Africa. In: Harlan JR, Wet JMJD, Stemler ABL, eds. Origins of African Plant Domestication. Berlin, New York: DE GRUYTER MOUTON, 465–478.

Harris K, Subudhi PK, Borrell A, Jordan D, Rosenow D, Nguyen H, Klein P, Klein R, Mullet J. 2007. Sorghum stay-green QTL individually reduce post-flowering drought-induced leaf senescence. Journal of Experimental Botany 58: 327–338.

Hayes CM, Weers BD, Thakran M, Burow G, Xin Z, Emendack Y, Burke JJ, Rooney WL, Mullet JE. 2016. Discovery of a Dhurrin QTL in Sorghum: Co-localization of Dhurrin Biosynthesis and a Novel Stay-green QTL. Crop Science 56: 104–112.

Hu Z, Olatoye MO, Marla S, Morris GP. 2019. An integrated genotyping-by-sequencing polymorphism map for over 10,000 sorghum genotypes. The Plant Genome 12: 1–15.

Korte A, Farlow A. 2013. The advantages and limitations of trait analysis with GWAS: a review. Plant Methods 9: 29.

Lasky JR, Upadhyaya HD, Ramu P, Deshpande S, Hash CT, Bonnette J, Juenger TE, Hyma K, Acharya C, Mitchell SE, et al.2015. Genome-environment associations in sorghum landraces predict adaptive traits. Science Advances 1: e1400218.

Leiser WL, Rattunde HF, Weltzien E, Cisse N, Abdou M, Diallo A, Tourè AO, Magalhaes JV, Haussmann BI. 2014. Two in one sweep: aluminum tolerance and grain yield in P-limited soils are associated to the same genomic region in West African Sorghum. BMC Plant Biology 14: 206.

Li D, Dossa K, Zhang Y, Wei X, Wang L, Zhang Y, Liu A, Zhou R, Zhang X. 2018a. GWAS Uncovers Differential Genetic Bases for Drought and Salt Tolerances in Sesame at the Germination Stage. Genes 9: 87.

Li X, Guo T, Mu Q, Li X, Yu J. 2018b. Genomic and environmental determinants and their interplay underlying phenotypic plasticity. Proceedings of the National Academy of Sciences 115: 6679–6684.

Lipka AE, Tian F, Wang Q, Peiffer J, Li M, Bradbury PJ, Gore MA, Buckler ES, Zhang Z. 2012. GAPIT: genome association and prediction integrated tool. Bioinformatics 28: 2397–2399.

Mace E, Innes D, Hunt C, Wang X, Tao Y, Baxter J, Hassall M, Hathorn A, Jordan D. 2019. The Sorghum QTL Atlas: a powerful tool for trait dissection, comparative genomics and crop improvement. Theoretical and Applied Genetics 132: 751–766.

Mace ES, Li Y, Prentis PJ, Bian L, Campbell BC, Hu W, Innes DJ, Han X, Cruickshank A, Dai C, et al.2013. Whole-genome sequencing reveals untapped genetic potential in Africa’s indigenous cereal crop sorghum. Nature Communications 4.

Mauboussin J-C, Gueye J, N’Diaye M. 1977. L’amélioration du sorgho au Sénégal. Agronomie Tropicale 32: 303–310.

McCormick RF, Truong SK, Sreedasyam A, Jenkins J, Shu S, Sims D, Kennedy M, Amirebrahimi M, Weers BD, McKinley B, et al.2018. The Sorghum bicolor reference genome: improved assembly, gene annotations, a transcriptome atlas, and signatures of genome organization. The Plant Journal 93: 338–354.

McCouch SR, Wright MH, Tung C-W, Maron LG, McNally KL, Fitzgerald M, Singh N, DeClerck G, Agosto-Perez F, Korniliev P, et al.2016. Open access resources for genome-wide association mapping in rice. Nature Communications 7: 1–14.

Mendiburu F. 2009. Agricolae: statistical procedures for agricultural research.

Morris GP, Ramu P, Deshpande SP, Hash CT, Shah T, Upadhyaya HD, Riera-Lizarazu O, Brown PJ, Acharya CB, Mitchell SE, et al.2013. Population genomic and genome-wide association studies of agroclimatic traits in sorghum. Proceedings of the National Academy of Sciences of the United States of America 110: 453–458.

Mullet J, Morishige D, McCormick R, Truong S, Hilley J, McKinley B, Anderson R, Olson SN, Rooney W. 2014. Energy Sorghum--a genetic model for the design of C4 grass bioenergy crops. Journal of Experimental Botany 65: 3479–3489.

Mundia CW, Secchi S, Akamani K, Wang G. 2019. A regional comparison of factors affecting global sorghum production: the case of north america, asia and africa’s sahel. Sustainability 11: 2135.

Murphy RL, Klein RR, Morishige DT, Brady JA, Rooney WL, Miller FR, Dugas DV, Klein PE, Mullet JE. 2011. Coincident light and clock regulation of pseudoresponse regulator protein 37 (PRR37) controls photoperiodic flowering in sorghum. Proceedings of the National Academy of Sciences 108: 16469–16474.

Murphy RL, Morishige DT, Brady JA, Rooney WL, Yang S, Klein PE, Mullet JE. 2014. Ghd7 (Ma6) represses sorghum flowering in long days: Ghd7 alleles enhance biomass accumulation and grain production. The Plant Genome 7: 1–10.

Paterson AH, Bowers JE, Bruggmann R, Dubchak I, Grimwood J, Gundlach H, Haberer G, Hellsten U, Mitros T, Poliakov A, et al.2009. The Sorghum bicolor genome and the diversification of grasses. Nature 457: 551–556.

Peterson BG, Carl P, Boudt K, Bennett R, Ulrich J, Eric Zivot, Lestel M, Balkissoon K, Wuertz D. 2014. Performanceanalytics: econometric tools for performance and risk analysis. version 1.4.4000 from r-forge.

R Core Team RC. 2016. A language and environment for statistical computing. R Foundation for statistical computing, 2015; Vienna, Austria.

Rao NN, Prasad K, Kumar PR, Vijayraghavan U. 2008. Distinct regulatory role for RFL, the rice LFY homolog, in determining flowering time and plant architecture. Proceedings of the National Academy of Sciences 105: 3646–3651.

Rodríguez-Álvarez MX, Boer MP, van Eeuwijk FA, Eilers PHC. 2018. Correcting for spatial heterogeneity in plant breeding experiments with P-splines. Spatial Statistics 23: 52–71.

Shin J-H, Blay S, McNeney B, Graham J. 2006. Ldheatmap: an r function for graphical display of pairwise linkage disequilibria between single nucleotide polymorphisms | shin | journal of statistical software. 16.

Tardieu F. 2012. Any trait or trait-related allele can confer drought tolerance: just design the right drought scenario. Journal of Experimental Botany 63: 25–31.

Traore SB, Ali A, Tinni SH, Samake M, Garba I, Maigari I, Alhassane A, Samba A, Diao MB, Atta S, et al.2014. AGRHYMET: A drought monitoring and capacity building center in the West Africa Region. Weather and Climate Extremes 3: 22–30.

Tuinstra MR, Grote EM, Goldsbrough PB, Ejeta G. 1996. Identification of quantitative trait loci associated with pre-flowering drought tolerance in sorghum. Crop Science 36: 1337–1344.

Tuinstra MR, Grote EM, Goldsbrough PB, Ejeta G. 1997. Genetic analysis of post-flowering drought tolerance and components of grain development in Sorghum bicolor (L.) Moench. Molecular Breeding 3: 439–448.

Upadhyaya HD, Dwivedi SL, Vetriventhan M, Krishnamurthy L, Singh SK. 2017. Post-flowering drought tolerance using managed stress trials, adjustment to flowering, and mini core collection in sorghum. Crop Science 57: 310–321.

Vadez V, Deshpande S, Kholova J, Ramu P, Hash CT. 2013. Molecular Breeding for Stay-Green: Progress and Challenges in Sorghum. In: Varshney RK, Tuberosa R, eds. Translational Genomics for Crop Breeding. Chichester, UK: John Wiley & Sons Ltd, 125–141.

Vadez V, Krishnamurthy L, Hash CT, Upadhyaya HD, Borrell AK. 2011. Yield, transpiration efficiency, and water-use variations and their interrelationships in the sorghum reference collection. Crop and Pasture Science 62: 645–655.

Vavilov NI. 2009. Origin and geography of cultivated plants (D Love, Tran.). Cambridge: Cambridge University Press.

Wang J, Hu Z, Upadhyaya HD, Morris GP. 2020. Genomic signatures of seed mass adaptation to global precipitation gradients in sorghum. Heredity 124: 108–121.

Wendorf F, Close AE, Schild R, Wasylikowa K, Housley RA, Harlan JR, Królik H. 1992. Saharan exploitation of plants 8,000 years BP. Nature 359: 721–724.

Wolabu TW, Zhang F, Niu L, Kalve S, Bhatnagar-Mathur P, Muszynski MG, Tadege M. 2016. Three FLOWERING LOCUS T-like genes function as potential florigens and mediate photoperiod response in sorghum. New Phytologist: 1–14.

Xu W, Subudhi PK, Crasta OR, Rosenow DT, Mullet JE, Nguyen HT. 2000. Molecular mapping of QTLs conferring stay-green in grain sorghum (Sorghum bicolor L. Moench). Genome 43: 461–469.

Yang S, Murphy RL, Morishige DT, Klein PE, Rooney WL, Mullet JE. 2014. Sorghum phytochrome B inhibits flowering in long days by activating expression of SbPRR37 and SbGHD7, repressors of SbEHD1, SbCN8 and SbCN12. PLoS ONE 9: e105352.

Yano K, Yamamoto E, Aya K, Takeuchi H, Lo P, Hu L, Yamasaki M, Yoshida S, Kitano H, Hirano K, et al.2016. Genome-wide association study using whole-genome sequencing rapidly identifies new genes influencing agronomic traits in rice. Nature Genetics 48: 927–934.

Yu J, Buckler ES. 2006. Genetic association mapping and genome organization of maize. Current Opinion in Biotechnology 17: 155–160.

Yuan Y, Xing H, Zeng W, Xu J, Mao L, Wang L, Feng W, Tao J, Wang H, Zhang H, et al.2019. Genome-wide association and differential expression analysis of salt tolerance in Gossypium hirsutum L at the germination stage. BMC Plant Biology 19: 394.

Zhao Y, Qiang C, Wang X, Chen Y, Deng J, Jiang C, Sun X, Chen H, Li J, Piao W, et al.2019. New alleles for chlorophyll content and stay-green traits revealed by a genome-wide association study in rice (Oryza sativa). Scientific Reports 9: 2541.

